# Mechanism of cooperative strigolactone perception by the MAX2 ubiquitin ligase–receptor–substrate complex

**DOI:** 10.1101/2024.11.17.623987

**Authors:** Alexandra I. Vancea, Brandon Huntington, Wieland Steinchen, Christos Savva, Umar F. Shahul Hameed, Stefan T. Arold

## Abstract

Strigolactones (SLs) are a group of plant hormones that regulate various aspects of plant growth and development. Additionally, SLs exuded into the soil promote symbiotic relationships with arbuscular mycorrhizal fungi and stimulate the germination of parasitic plants such as *Striga hermonthica*. The binding and hydrolysis of SLs by their receptors (D14 in Arabidopsis and HTL in *Striga*) promote the ubiquitination of transcriptional repressors by Skp1–cullin–F-box (SCF)–type E3 ubiquitin ligases. The mechanistic link between SL perception by D14/HTL and substrate recognition by the E3 remains unclear. We identified an E3–HTL–substrate complex that is sufficiently stable for cryogenic electron microscopy. This complex, composed of SKP1 (ASK1) and substrate (SMAX1) from Arabidopsis, and Striga F-box (MAX2) and SL receptor (HTL7), reveals that the substrate engages in a bidentate association through its N and D2 domains. This interaction, which is both conformationally and compositionally dynamic, directly and allosterically stabilises the MAX2–(SL)HTL7 complex and affects the positioning of ASK1 relative to MAX2. This dynamic positioning influences the proximity between the substrate D2 domain and the ubiquitin-conjugated E2 enzyme. This work advances our understanding of how E3 ligases in plants translate hormone perception into genetic adaptations.

## INTRODUCTION

Eukaryotes utilise a cascade of enzymes, known as E1, E2, and E3, to post-translationally modify substrate proteins with ubiquitin^1–3^. Ubiquitination often leads to proteasome-mediated degradation of the substrate but can also result in altered location or activity^4–6^. Thus, ubiquitination is essential for maintaining a healthy proteome and enables cells to rapidly adapt to changing conditions. In plants, the genetic diversity of the E3 ubiquitin ligase family has expanded substantially, with more than 1,000 potential members, highlighting their increased importance for signalling and adaptation in these sessile organisms^7–9^.

E3 ligases mediate the ubiquitination of substrates by linking them with ubiquitin-conjugated E2 proteins. A major family of E3 ligases in both plants and animals is the multisubunit SCF complex, named after its canonical core components Skp1, cullin, and F-box proteins^10–13^. In these complexes, cullins bind to ubiquitin-loaded E2s (and associated RING-box [Rbx] protein), F-box proteins bind to the substrates, and Skp1 connects both parts.

Reflecting their role in recognizing diverse substrates, nearly 900 different F-box proteins have been identified in plants^14–16^. Whereas animal F-box proteins primarily recognize post-translationally modified substrates^17^, recent findings suggest that plant F-box proteins have evolved sophisticated recognition mechanisms to also function as environmental sensors^7,18^. These sensors link phytohormone perception with proteomic changes. Initial structural analyses have revealed how SCF E3 ligases mediate the perception of plant hormones such as auxin and jasmonate^18–20^. In both cases, the hormone associates with the leucine-rich repeat (LRR) domain of the F-box protein, creating a shared binding site for the substrates. The subsequent ubiquitination and degradation of these substrates, which are transcriptional repressors, lead to changes in gene expression.

The perception of strigolactones (SLs) also involves SCF E3 ligases, but with additional layers of complexity. Strigolactones are plant hormones that play diverse roles in regulating plant architecture, adaptation, defence, and plant–rhizospheric communications^21,22^. SLs are carotenoid derivatives characterised by a butenolide ring (D-ring) linked by an enol ether bridge to a tricyclic lactone (ABC-ring) in canonical SLs, or to a different moiety in non-canonical SLs^23–25^. Unlike auxin and jasmonate, SLs do not bind directly to their F-box protein (MORE AXILLARY GROWTH 2 (MAX2) in Arabidopsis and DWARF3 (D3) in rice) but to DWARF14 (D14)^26,27^. D14, derived from an α/β-hydrolase, retains low hydrolase activity allowing cleavage of the SL enol ether bridge while retaining the D-ring^28,29^. SL binding and cleavage facilitate structural changes in D14, enabling it to associate with MAX2^28,30^. The F-box–D14 complex then binds to substrates, which are the transcriptional repressors DWARF53 (in rice) or SUPPRESSOR OF MAX2 1 (SMAX1)– like (SMXL) 6, 7, and 8 (in Arabidopsis)^31–35^. These substrates all share a common domain structure composed of an N-terminal Double Clp-N motif (N), a first ATPase domain (D1), a middle domain (M), and a C-terminal second ATPase domain (D2). According to structural predictions, these domains are flexibly connected by linkers that vary in sequence and length.

SLs exuded from the roots of host plants also trigger the germination of the parasitic plant *Striga hermonthica*^36^. Once germinated, *S. hermonthica* attaches to the roots of host plants and starts syphoning off their nutrients. This parasitism leads to significant crop losses, amounting to billions of dollars annually, and exacerbates food insecurity in affected regions^37–39^. *S. hermonthica* has adapted the SCF–D14 SL recognition mechanism by utilising a family of D14 homologues known as HYPOSENSITIVE TO LIGHT (HTL) proteins^40^. These HTLs link the F-box protein *Sh*MAX2 to the transcriptional repressors responsible for seed germination, presumably Striga paralogues of the SMAX1. Among *S. hermonthica*’s HTLs, *Sh*HTL7 is the most sensitive to SLs^41–43^. Efforts to design drugs to control *S. hermonthica* germination have been challenging due to the homology of the SL perception mechanisms of the parasite and its host plants^39,44–46^.

Although there is general agreement on the SLs response mechanism in plants, several key aspects remain elusive and controversial. These include the nature of the SL degradation product bound to D14/HTL during its association with MAX2/D3^47,48^, and the mechanism by which the association of D14/HTL with MAX2/D3 leads to substrate recognition and ubiquitination^49,50^. These knowledge gaps are primarily due to the lack of structural information on the complex formed between the F-box protein, D14/HTL, and the substrate. While accumulating evidence suggests that the substrate itself contributes to the stabilisation and specificity of the SCF–D14/HTL–substrate complex, only structures of Skp1–F-box alone or Skp1– F-box–D14 have been reported^28,51,52^. Notably, a recent attempt to obtain the SCF–D14/HTL– substrate complex using cryo-electron microscopy (cryo-EM) failed to capture density for the D53 substrate, even though D53 was necessary to stabilise the MAX2–D14 interaction^53^. This presents a conundrum: the substrate stabilises the complex and is subsequently ubiquitinated, yet the interactions are so transient that they cannot be captured by structural methods.

In this study, we exploit species-specific idiosyncrasies to identify an active signalling complex that is sufficiently stable to reveal the substrate using carefully timed cryogenic grid preparation for cryo-EM analysis. This complex, composed of Arabidopsis SKP1 (ASK1), *Sh*MAX2, *Sh*HTL7, and *At*SMAX1, demonstrates how a bidentate and markedly dynamic interaction with *At*SMAX1 produces the stability and specificity of the SCF^MAX2^–HTL complex, while allosterically affecting the positioning of the substrate toward the E2 enzyme. Our findings reveal a new mechanism for SCF-mediated phytohormone response and allow us to rationalise previous biochemical and functional observations.

## RESULTS

### IDENTIFICATION OF STABLE COMPLEXES

The interaction between D14/HTLs and the F-box proteins requires the slow hydrolysis of SLs by the receptor and is only stable for a few hours *in vitro*^26,29,54^. To capture this slow-onset transient complex, we used single-particle analysis (SPA) with cryogenic electron microscopy (cryo-EM) to image samples vitrified within a 2-3 h time window after addition of GR24, a synthetic SL analogue. Complexes were reconstituted by combining D14/HTL receptors produced in *Escherichia coli* with SKP1 and F-box proteins co-expressed in insect cells (see **Methods**). Our initial cryo-EM structural analysis focussed on Striga HTL7, rice D3, and Arabidopsis ASK1 as this complex showed GR24-dependent stable binding on size-exclusion chromatography (SEC) and pull-down studies (**Supplementary Fig. 1A-C**). Cryo-EM grids showed severe preferred particle orientation which we addressed by tilting the stage. Nonetheless, these grids only yielded the structure of the *At*ASK1–*Os*D3 complex to a maximum resolution of 4.3 Å (**Fig. 1A, Supplementary Fig. 1F-I**), without the SL-bound *Sh*HTL7 receptor, suggesting that the tripartite complex was too unstable under the cryogenic preparation conditions used. Attempts to stabilise the complex through chemical cross-linking were unsuccessful.

**Fig. 1:**
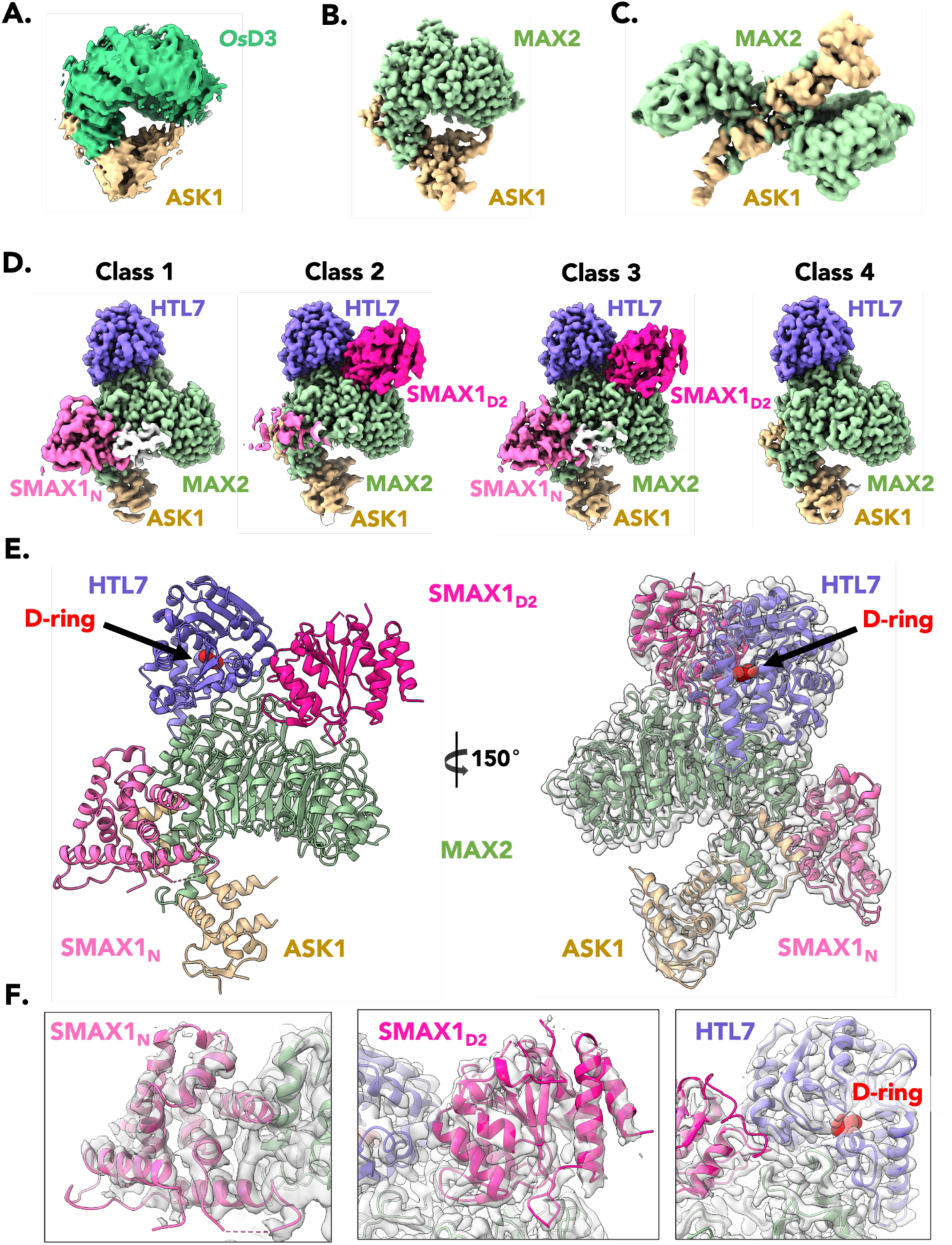
Cryo-EM structures of the ASK1–F-box–receptor–substrate complexes. **A.** Cryo-EM density of ASK1–*Os*D3 in the presence of HTL7 and the strigolactone analogue GR24 (**EMD-62415**). ASK1 and *Os*D3 are coloured yellow and green, respectively. Cryo-EM density of HTL7 was not visible. **B-D.** Compositional assemblies of ASK1–MAX2–HTL7–SMAX1 in the presence of GR24. ASK1, MAX2, and HTL7 are coloured yellow, green and purple, respectively. SMAX1_N_ is coloured light pink and SMAX1_D2_ is coloured dark pink. Unassigned cryo-EM density is coloured white. **B.** Cryo-EM density of apo-ASK1–MAX2 (**EMD-62408**). **C.** Cryo-EM density of the 2:2 ASK1– MAX2 dimer (**EMD-62414**). **D.** Cryo-EM density of ASK1–MAX2–HTL7 classes. Class 3 is a sub-class of Class 2, following masked classification around SMAX1_N_. (Class 1, **EMD-62397**; Class 2, **EMD-62401**; Class 3, **EMD-62417**; Class 4, **EMD-62407**). **E.** Complete atomic model of ASK1– MAX2–HTL7–SMAX1 from Class 3 (**PDB 9KLV**), coloured as in B-D. Cryo-EM density of Class 3 is shown as a transparent grey surface. The GR24 D-ring atoms are shown as red spheres and highlighted by black arrows. **F.** Magnified view of the fit of SMAX1_N_ in the cryo-EM density of Class 1 (left), and SMAX1_D2_ (middle) and HTL7 (right) in the cryo-EM density of Class 2. Cryo-EM densities are shown as transparent grey surfaces.

Next, based on literature reports^53,55^, we included the ubiquitination substrate to stabilise the complex for cryo-EM. We selected *At*SMAX1 as the substrate because it has been previously reported to interact with *Sh*HTL7 and both Striga and Arabidopsis MAX2^55,56^, and because reliable Striga SMAX1 orthologues were unavailable (our work started before the annotated Striga proteome was reported^57^. SEC and pull-down experiments indicated that the most stable complex in the presence of GR24 was formed by HTL7 and MAX2 from *S. hermonthica*, along with *A. thaliana* ASK1 and SMAX1 (**Fig. 2A,B**). Conversely, *At*SMAX1 interacted only weakly with *Os*D3– *At*ASK1–*Sh*HTL7 in the presence of GR24 (**Supplementary Fig. 1J**). We therefore focussed on *At*ASK1, *Sh*MAX2, *Sh*HTL7, and *At*SMAX1, which we hitherto refer to simply as ASK1, MAX2, HTL7, and SMAX1, respectively, unless disambiguation is required.

### STRUCTURAL LANDSCAPE OF THE SCF^MAX2^–RECEPTOR–SUBSTRATE COMPLEXES

For cryo-EM analysis, purified ASK1, MAX2, HTL7, and SMAX1 were mixed and incubated with GR24. Samples were run through SEC, and the peak fraction was concentrated and used for cryo-EM grids. SPA of these grids yielded three compositional assemblies: (1) ASK1–MAX2; (2) a 2:2 ASK1–MAX2 dimer; and (3) the ASK1–MAX2–HTL7 complex, each of them representing approximately one third of the curated particles (**Supplementary Fig. 2**). For ASK1–MAX2, the obtained 524k particles produced a 2.7 Å resolution structure (**Fig. 1B, Supplementary Fig. 3**). Although we reconstructed the 2:2 ASK1–MAX2 dimer using 309k particles, atomic resolution modelling was precluded by high anisotropy (see below) (**Fig. 1C, Supplementary Fig. 4**).

**Fig. 2:**
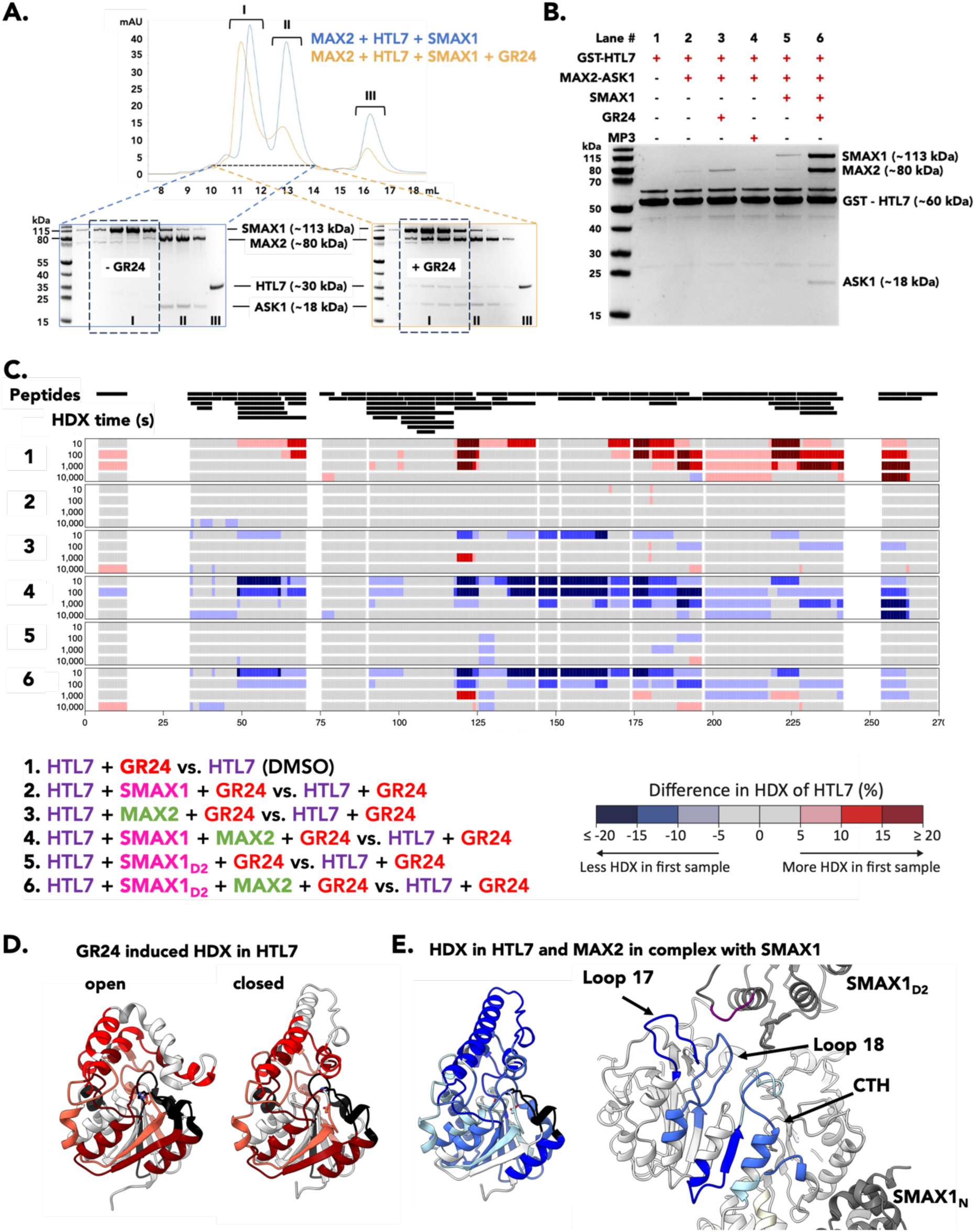
Biochemical confirmation of ASK1–MAX2–HTL7–SMAX1 interactions. **A.** SEC profiles (top) and corresponding Coomassie-stained SDS-PAGE (bottom) of ASK1–MAX2–HTL7–SMAX1 with (orange; right) and without (blue; left) the addition of GR24. Solid lines show the absorbance at 280 nm. Roman numerals indicate the peak fractions analysed. The approximate molecular weight of each protein is labelled next to the reference ladder. The dotted black box highlights the fractions containing the entire complex. **B.** Coomassie stained SDS-PAGE of GST pull-down assays using GST-HTL7 as the bait and combinations of ASK1–MAX2, SMAX1, GR24, and MP3 as targets. Plus signs denote presence and minus signs absence of species in each pull-down assay. Approximate molecular weights are labelled. **C.** Each black bar represents a peptide of HTL7 identified in HDX-MS experiments. The difference in HDX of HTL7 of pairwise comparisons between HTL7-containing samples is displayed on its amino acid sequence. Different tones of red and blue indicate HTL7 residues that incorporate more or less deuterium, respectively, in the first state of the pairwise comparison **D.** Deuterium enriched peptides mapped on the open (PDB 5Z8P) and closed structure of HTL7 from this study (**PDB 9KKX**). **E.** Protected regions of HTL7+GR24 and MAX2 in complex with SMAX1 mapped on the structure (**PDB 9KLV**). The black coloured regions had no peptide coverage.

The ASK1–MAX2–HTL7 structures showed the same seahorse-shaped topology as observed previously in the crystal structure of the ASK1–*Os*D3–*At*D14 complex (PDB 5HZG)^28^, with HTL7 as the head part, the solenoid-shaped leucine-rich repeat (LRR) domain of MAX2 as the body, and the N terminal F-box motif of MAX2 together with ASK1 as the tail (**Fig. 1D,E**).

Further 3D classification of the ASK1–MAX2–HTL7 assembly identified different additional densities that represented domains of SMAX1 (**Supplementary Fig. 2**). Class 1 (with 60k particles, refined to 2.8 Å resolution) contained additional density docked on the leucine-rich repeat (LRR) domain of MAX2 in proximity to ASK1 (**Fig. 1D**). Based on secondary structure features and AlphaFold structure predictions, we identified this density as the SMAX1 N domain (residues 10-167) (**Fig. 1E,F**). Class 2 (with 80k particles refined to 2.9 Å resolution) showed weaker density for the N domain, but featured a clear additional density between HTL7 and MAX2, which we identified as the SMAX1 D2 domain (**Fig. 1D,E,F**). This D2 position suggested that it stabilises the interaction between HTL7 and MAX2. The observation that Class 1 and Class 2 showed good density for either the N or the D2 domain of SMAX1 raised the question of whether both domains can bind to MAX2 simultaneously. Further classification of Class 2 identified a subset of 17k particles (termed Class 3) that produced a structure with density for both N and D2 domains, but at a lower resolution of 3.3 Å (**Fig. 1D**). A final Class 4 contained the complex that only showed density for ASK1, MAX2 and HTL7 (44k particles, 2.9 Å) (**Fig. 1D).** However, in these structures, the C186 loop of LRR4 was in the position likely associated with the presence of SMAX1 (see below). Indeed, the MAX2 LRR domains across classes 1-4 did not show significant changes (RMSD < 0.5 Å).

Collectively, these structures suggested that the N and D2 domains of SMAX1 bind to the ASK1–MAX2–HTL7 complex with a significant flexibility (conformational heterogeneity), where focussed refinement results in different degrees of visibility of the N and D2 domain density. However, our data also supports the existence of compositional heterogeneous complexes, where either N or D2 are bound.

### BIOCHEMICAL CONFIRMATION OF THE COMPLEX ARCHITECTURE

Our cryo-EM analysis provided several observations: (i) HTL7 undergoes large structural rearrangements in its lid domain upon binding to MAX2, (ii) SMAX1 binds to the complex with at least two domains (N and D2) in a flexible and dynamic manner, (iii) D2 binding stabilises the association of HTL7 with MAX2, and (iv) the N domain established only few direct interactions with ASK1. To independently validate these findings in solution, we employed hydrogen-deuterium exchange coupled with mass spectrometry (HDX-MS).

We compared the changes in HTL7 when complexed with ASK1–MAX2 alone, and in combination with full-length SMAX1 and the D2 domain only, all in the presence of GR24. We obtained a high peptide coverage for all the components (95% for ASK1, 97% for MAX2, 78% for HTL7, and 90% for SMAX1) enabling us to assess the dynamics of all components **(Fig. 2C; Supplementary Fig. 5; Supplementary Fig. 6; Supplementary Dataset 1**).

The presence of the SL GR24 induced strong HDX deprotection of HTL7 compared to the DMSO mock (**Fig. 2C,D; Supplementary Fig. 5A,B**). This deprotection indicated increased malleability of the SL-bound state, consistent with previously reported thermal destabilisation of D14 and HTL7 in the presence of SLs^29,30^. HDX of GR24-HTL7 was reduced by MAX2 alone. This HDX reduction was increased in presence of SMAX1_D2_ and even more so when full-length SMAX1 was present, confirming that the D2 domain stabilises GR24-HTL7 on MAX2 (**Fig. 2C,E; Supplementary Fig. 5A,B**). Correspondingly, HDX of MAX2 was reduced in the presence of GR24-HTL7. This reduction was amplified in the presence of SMAX1 and to a lesser degree by D2 alone (**Supplementary Fig. 5C**,D). Although SMAX1 had a clear effect on the HDX of HTL7 and MAX2, we did not observe significant changes in the HDX of SMAX1 (full-length or D2) in the presence of the other components, in line with a transient and dynamic interaction (**Supplementary Fig. 6A,B**). Finally, consistent with the structural analysis, ASK1 did not show HDX changes in any of the complexes tested (**Supplementary Fig. 6C,D**).

Additionally, MAX2 and GST-HTL7 bound only weakly in GST pull-down assays in presence of GR24, and did not co-elute in SEC in the absence of SMAX1 (**Fig. 2A; Supplementary Fig. 7A**). The presence of SMAX1 increased GR24-induced co-precipitation and co-elution of MAX2 and HTL7, further confirming the stabilising role of SMAX1 (**Fig. 2A,B)**. However, also the presence of SMXL6 and SMXL7, considered to be the D14-specific substrates, allowed GST-HTL7 to co-precipitate MAX2 in a GR24-dependent manner (**Supplementary Fig. 7B**). Conversely, the homologous complexes *Os*D3–HTL7 or *At*MAX2–HTL7 showed an association on SEC even without SMAX1 homologues (**Supplementary Fig. 1A-C; Supplementary Fig. 7C**). We also observed that the non-canonical SL MP3^58^ was markedly less potent in inducing the MAX2–HTL7 and *Os*D3–HTL7 complex (**Fig. 2B; Supplementary Fig. 1A-C**). Thus, the E3-target interactions appear to vary with paralogues and hormone isoforms.

### THE CLOSED D-RING IS BOUND TO THE CATALYTIC HIS IN THE SIGNALLING STATE

In our cryo-EM structures, HTL7 is attached to the C-terminal LRRs 17-20 of MAX2, with its lid domain in the closed conformation (**Fig. 1E,F; Fig. 3A**). This position and conformation of HTL are consistent with those reported for *At*D14 in its crystallographic complex with ASK1–*Os*D3 (PDB 5HZG)^28^ (**Supplementary Fig. 8A**). HTL7 clearly displays an additional density attached to the catalytic histidine H246 (**Fig. 3B**). This density is in agreement with a covalently bound closed D-ring (termed the C_5_H_5_O_2_ modification)^43,56,59^, but not with the open CLIM linked to both S95 and H246, as proposed for the ASK1–*Os*D3–*At*D14 crystal structure (PDB 5HZG)(**Supplementary Fig. 8B**)^28^. Our cryo-EM structure was captured within the hour-long SL signalling window, suggesting that the closed D-ring corresponds to the signalling state, in agreement with published mass spectrometry results of HTL7 incubated with GR24^43,56^. The D-ring is tightly encapsulated by hydrophobic interactions involving residues Y26, M96, Y174, L178, and I193. These residues are distant from the active site residues in the open form of HTL, but close in on the active site in the closed MAX2-bound HTL conformation, leaving no space for the ABC rings of an uncleaved SL (**Fig. 3C**). Jointly with our HDX data, this observation clarifies that SL cleavage must occur before the docking of HTL7 onto the MAX2 LRR surface can be completed.

**Fig. 3:**
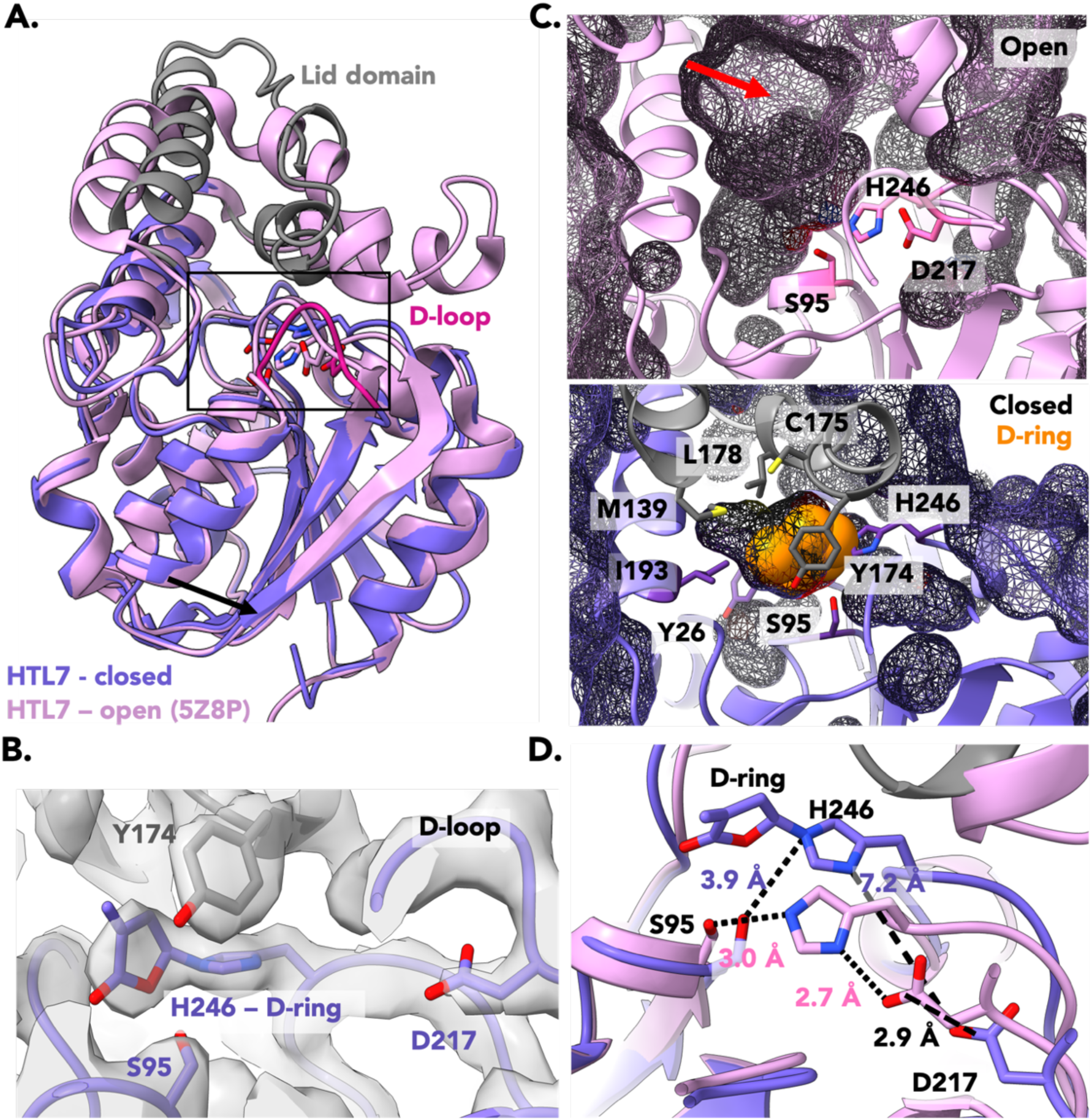
MAX2–HTL7 is closed with the GR24 D-ring trapped in its active site. **A.** Superimposition of HTL7 from our cryo-EM structure (**PDB 9KKX**; purple) and the open-conformation crystal structure of HTL7 (5Z8P; pink). The collapsed lid domain of our model is coloured grey and the D-loop is coloured red. The catalytic triad residues are shown as sticks. The black box outlines the active site and magnified views in the following panels. **B**. Fit of the D-ring covalently attached to H246 (H246–D-ring) in the cryo-EM density. Catalytic triad residues are shown as sticks and labelled. **C.** Hydrophobic enclosure (mesh outline) in the open (top) and our closed (bottom) HTL7 active site. The GR24 D-ring is shown as orange spheres. Red arrows highlight the active site entrance. The collapsed lid domain is coloured grey. Catalytic triad residues are shown as sticks and labelled (top). Residues that form the hydrophobic enclosure in the closed conformation are shown as sticks and labelled (bottom). **D.** Superimposition of catalytic triad residues in closed and open conformations. Distances between atoms are shown as dotted lines and labelled.

In the MAX2-bound HTL7, the loop containing H246 is shifted away from S95 to a distance of 3.9 Å, compared to 3.0 Å in the open state, likely to accommodate the covalently attached D-ring. The catalytic D217, located on the so-called D-loop, is also displaced by 2.9 Å relative to open HTL7. The combined movement of H246 and D217 results in a distance of more than 7 Å between their side chains, compared to 2.7 Å in the open form (**Fig. 3D**) (PDB 5Z8P)^60^, disrupting the catalytic triad. Along with the tight hydrophobic enclosure that prevents nucleophilic attack by a water molecule, this architecture stabilises the covalently bound D-ring and the resulting signalling MAX2 complex.

In the crystal structure of *Os*D3–*At*D14, the D-loop was not resolved and was proposed to move out to associate with D53/SMXLs^29^. In our structure, the D-loop was well resolved and positioned more than 10 Å away from the SMAX1 D2 domain (**Supplementary Fig. 8C**). No unattributed density was visible near the D-loop, but we cannot exclude transient interactions with SMAX1 regions other than the N or D2 domains.

### STABILISATION OF THE MAX2–HTL7 INTERACTION BY THE SUBSTRATE

The SMAX1 D2 domain is required for substrate ubiquitination and degradation^61^. The closest sequence homologues for which an experimental structure exists are the AAA+ disaggregation ClpB proteins, which share up to 23% sequence identity. Our cryo-EM maps clearly resolve the D2 α/β core domain (residues 607–767, corresponding to the structured regions of the D2a annotation^61^) and the C-terminal region of D2b (residues 860–885) which adds a helix-strand element to the D2a α/β core **(Fig. 1F; Fig. 4A)**. Density for the remaining D2b domain residues 774-798 and 886-990 (forming a four-helix structure capped by a two-stranded β-sheet) only appears at lower thresholds in the cryo-EM map, indicating that this domain is more flexible (**Supplementary Fig. 9A**).

**Fig. 4:**
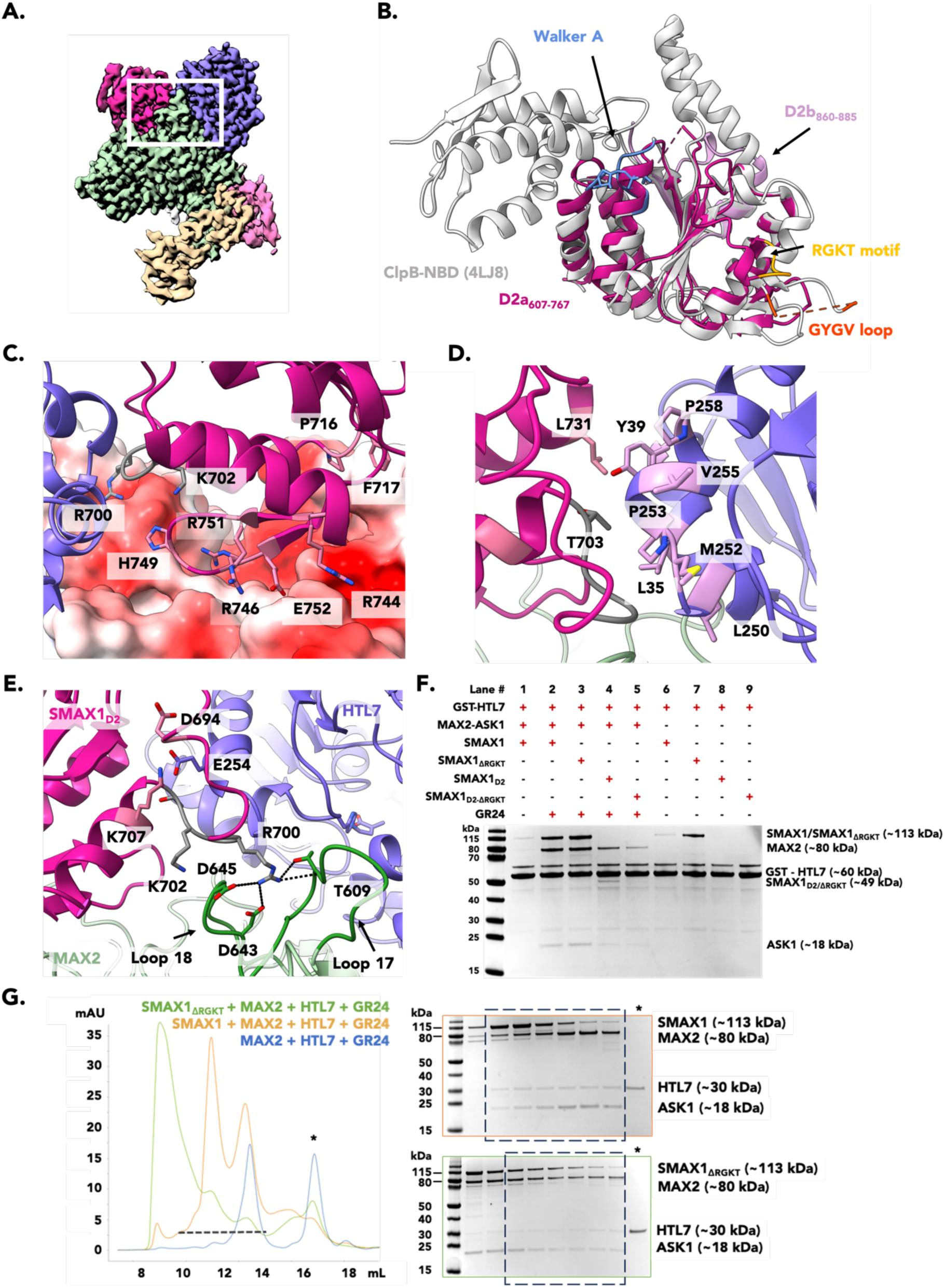
SMAX1_D2_ stabilises MAX2–HTL7. **A.** Reference view of ASK1–MAX2–HTL7–SMAX1 (**EMD-62417**) with a white box outlining the region magnified in panels B-E. **PDB 9KLD** is used for panels B-E. Proteins and domains are coloured as in Fig 1. **B**. SMAX1_D2a_ (residues 607-767; dark pink) and SMAX1_D2b_ (residues 607-767; light pink) superimposed on the nucleotide binding domain (NBD) of ClpB (4LJ8; white). The GYGV loop of ClpB (orange) and R^700^GKT motif of SMAX1_D2_ (yellow) are highlighted. The bound ADP shown as sticks and the Walker A fold of ClpB are coloured blue. The Walker A fold is not present in SMAX1_D2_. **C.** Polar interface of SMAX1_D2_–MAX2. MAX2 is depicted as a surface coloured by electrostatic potential from red (negatively charged) to blue (positively charged). Interacting residues of SMAX1_D2_ are shown as light pink sticks and labelled. The RGKT motif residues of SMAX1_D2_ are coloured grey and labelled. **D.** Hydrophobic interface of HTL7– SMAX1_D2_. The R^700^GKT motif residues of SMAX1_D2_ are coloured grey and labelled. Hydrophobic residues are shown as sticks and labelled. **E.** Interaction of MAX2–HTL7–SMAX1_D2_ through R700 of the SMAX1_D2_ R^700^GKT motif. The R^700^GKT motif residues of SMAX1_D2_ are coloured grey and labelled. LRR loops 17 and 18 of MAX2 are indicated with black arrows and hydrogen bonds and salt bridges with R700 are shown with dotted lines. Interacting residues are shown as sticks and labelled. **F.** Coomassie stained SDS-PAGE of GST pull-down assays using GST-HTL7 as a bait and combinations of ASK1–MAX2, full-length SMAX1, full-length SMAX1_ΔRGKT_, SMAX1_D2_, SMAX1_D2ΔRGKT_, and GR24. Plus signs denote presence and minus signs absence of species in each pull-down assay. Approximate molecular weights are labelled. **G.** SEC profiles (left) and corresponding Coomassie stained SDS-PAGE (right) of ASK1–MAX2–HTL7–SMAX1 and GR24 with (orange) and without (green) the SMAX1 RGKT motif versus MAX2–HTL7 and GR24 alone (blue). Solid lines show the absorbance at 280 nm. Asterisk and black dashed line and boxes highlight the fractions analysed by SDS-PAGE. The corresponding SDS-PAGE gel for each SEC profile is outlined with the same colour. The corresponding SDS-PAGE gel for MAX2–HTL7 and GR24 alone can be found in **Supplementary Fig. 7A**.

Structure superimposition of SMAX1_D2_ with the NBD2 of ClpB from *T. thermophilus* (PDB 4LJ8)^62^ produces an RMSD of 1.3 Å for the conserved α/β core (85 residues), but of more than 6 Å for all 166 residues (**Fig. 4A**). Indeed, SMAX1_D2_ differs in key functional elements, showing its structure and function have significantly derived from NBDs of hexameric AAA+ ATPases. We observed no evidence of a bound nucleotide in our structure, in agreement with the degenerate Walker A motif forming a helix that occludes the canonical nucleotide binding site (**Supplementary Fig. 9B,C**). The GYGV motif, which forms a pore loop in the ClpB hexamer, is replaced by the R^700^GKT motif in SMAX1_D2_ (**Fig. 4A; Supplementary Fig. 9C**). This motif is conserved among SMAX1 homologues **(Supplementary Fig. 9D**)^61,61^. Its deletion in SMAX1, D53, SMXL6, and SMXL7 abrogates GR24-induced degradation of these proteins *in planta*, suggesting that this motif is either ubiquitinated or stabilising the SCF–D14/HTL complex^32,34,35,61^.

In our cryo-EM model of the E3–substrate complex, D2 associates with an LRR surface formed through mostly charged and polar contacts (D2 residues R700, K702, R744, R746, H749, R751, and E752) with only two hydrophobic residues making significant contacts (P716 and F717) (**Fig. 4C,D**). This interaction buries 928 Å^2^ of solvent exposed surface. Conversely, the D2 contact surface with HTL7 is smaller (430 Å^2^ buried in total) and more hydrophobic, formed mostly by D2 residues G701, T703, and L731, engaging HTL7 residues L35, P39, Y39, L250, M252, P253, V255, and P258) (**Fig. 4E**) Additionally, HTL7 E254 is reaching into the D2 loop formed by residues 694 to 703, where E254 forms a hydrogen bond with the backbone carbonyl of D694, and an ionic interaction with K707.

The D2 residue K702 is mostly buried in the complex, making it an unlikely substrate for ubiquitination (**Fig. 4C,D**). However, D2 R700 stabilises the association between HTL7 and MAX2 by wedging between the MAX2 loops of LRRs 17 and 18 (residues 604–616 and 642–650, respectively) (**Fig. 4D).** In this position, R700 forms ionic bonds with MAX2 D643 and D645, and hydrogen bonds with T609. The LRR17 and LRR18 loops are in a similar position in the crystallographic ASK1–*Os*D3–*At*D14 complex (PDB 5HZG, where SMAX1 was absent), but the LRR17 loop was arranged differently in our *At*ASK1–*Sh*MAX2 apo structure (**Supplementary Fig. 8E**), indicating that D2 stabilises the D14/HTL-bound conformation of the LRR17 loop.

To probe the stabilising role of the R^700^GKT motif, we created RGKT deletion mutants in both full-length SMAX1 and SMAX1_D2_. In SEC and pull-down experiments, we observed that the ability to stabilise the ASK1–MAX2–HTL7–SMAX1 complex was reduced in the ΔRGKT mutants (**Fig. 4E,F**). This reduction was more pronounced in the D2 construct compared to full-length SMAX1 (**Fig. 4F**), consistent with the observation that D2 alone increased GR24-induced complex coprecipitation less than full-length SMAX1, due to the additional binding contributions from the N domain in the latter. These data support that the R^700^GKT motif is conserved for its role in allosterically stabilising the HTL7–MAX2 complex, rather than serving as a ubiquitination site. Taken together, these findings demonstrate that SMAX1_D2_ is an ATPase domain repurposed to function as a non-catalytic binding domain that helps stabilise the MAX2–HTL–substrate complex.

### IDIOSYNCRASIES IN THE INTERFACE BETWEEN MAX2 AND THE N DOMAIN

The SMAX1_N_ domain belongs to the Clp repeat (ClpR) family^63,64^. Accordingly, its two four-helix structure repeats (1: residues 12-83; and 2: residue 97-167) are related by a pseudo-twofold symmetry^65^. Structural superimposition with the ClpC1 N-terminal domain (NTD) (PDB 6PBA; sequence identity 17.3%)^66^ yielded an RMSD of 1 Å across the conserved helical structures and up to 5 Å across all pairs (**Fig. 5A, Supplementary Fig. 10A**).

**Fig. 5:**
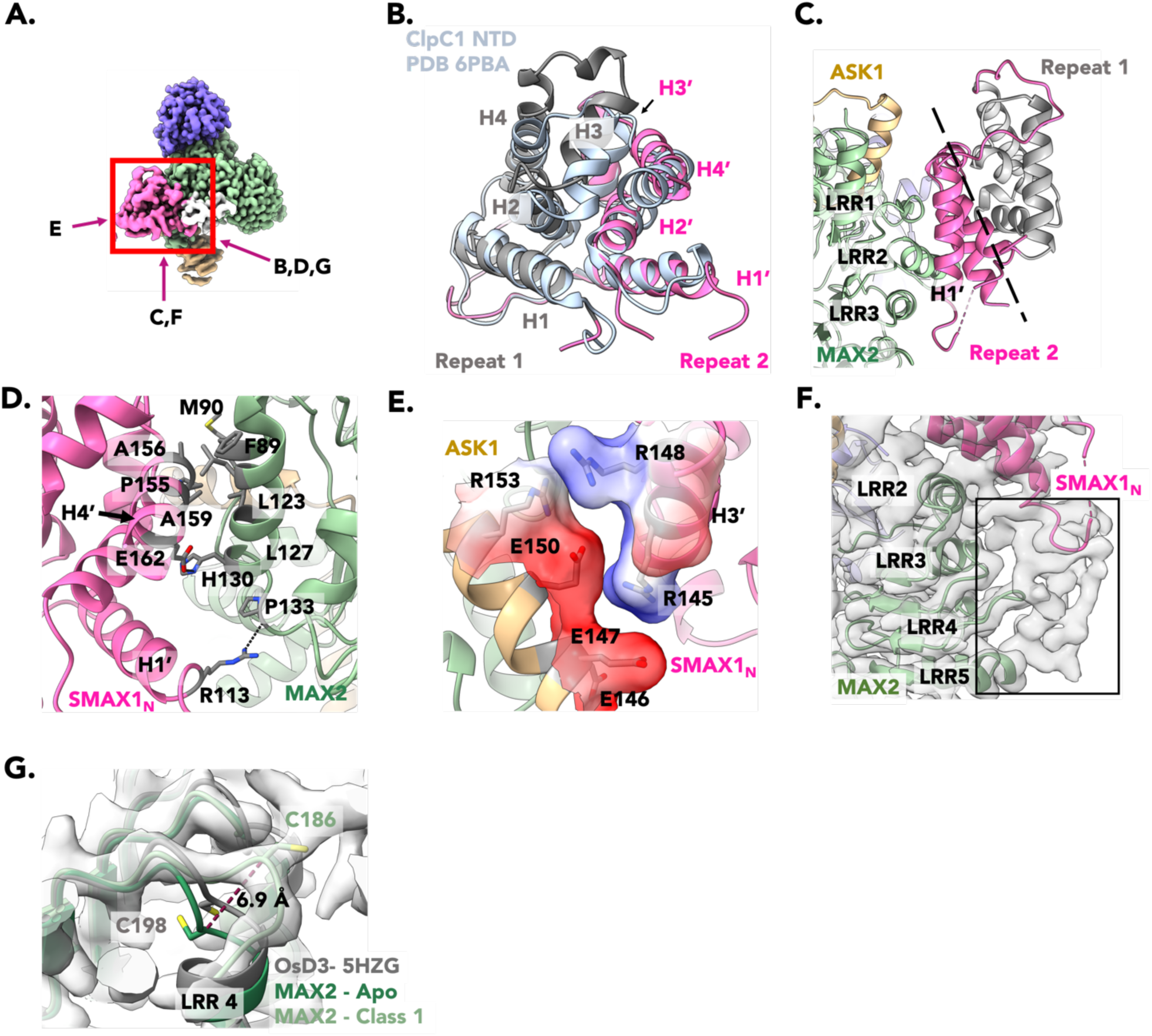
The MAX2–SMAX1_N_ interface is variable. **A.** Reference view of ASK1–MAX2–HTL7–SMAX1 (**EMD-62397**) with a red box outlining the region magnified in panels B-G. Arrows show the relative orientation of the corresponding panel. **PDB 9KKX** is used for panels B-G. Proteins and domains are coloured as in Fig 1. **B.** Repeat 1 (grey) and Repeat 2 (pink) of SMAX1_N_ superimposed on the N-terminal domain (NTD) of ClpC1 (6PBA; light blue). SMAX1_N_ helices are labelled, with prime marking the pseudosymmetric helix in Repeat 2. **C.** Interface of MAX2–SMAX1_N_ through Repeat 2. The pseudosymmetric axis of SMAX1_N_ is shown as a dotted line. The interacting LRR1-3 of MAX2 and H1’ of SMAX1_N_ are labelled. **D.** Hydrophobic interface of MAX2–SMAX1_N_. Interface residues are coloured grey, shown as sticks, and labelled. Additional hydrogen bonds between SMAX1_N_ residues E162 and R113 with MAX2 H130 and the carbonyl oxygen of P133 are shown as dotted lines. **E.** Electrostatic interface of ASK1–SMAX1_N_. Red represents negatively and blue represents positively charged surfaces. Interface residues are coloured grey, shown as sticks, and labelled. **F.** Unassigned cryo-EM density (black box) adjacent to SMAX1_N_ and LRR3-5 of MAX2. Cryo-EM density is shown as a transparent grey surface. **G.** Position of MAX2 C186 in ASK1– MAX2–HTL7–SMAX1 (**PDB 9KKX**; light green), apo-ASK1–MAX2 (**PDB 9KLL**; dark green) and *Os*D3 (PDB 5HZG; grey). Cryo-EM density of Class 1 (**EMD-62397**) is shown as a transparent grey surface. C198 of *Os*D3 and C186 of MAX2 are shown as sticks and labelled. The distance between C186 in apo-ASK1–MAX2 and Class 1 is shown as a dotted line.

Our structures show that repeat 2 of SMAX1_N_ binds to MAX2 LRRs 1 and 2, close to the ASK1 interface, burying 558 Å^2^ of surface area (**Fig. 5B)**. The hydrophobic side of the C-terminal SMAX1_N_ helix (H4’, residues S153 to N166) contacts a hydrophobic patch on the first and second LRR (residues M90, F89, L123, and L127) (**Fig. 5C**). This interaction is reinforced by additional hydrogen bonds and charge-charge interactions between R145 and R148 of the SMAX1_N_ H3’ helix and E146, E147, and E150 of ASK1 (**Fig. 5D**). An additional density, which we could not attribute to any protein region, is visible between the SMAX1_N_ H1’–H2’ loop and the loops connecting the helices of LRR 3 to 5 with their following β strands (**Fig. 5E**). In this complex, the helix–β loop (residues 183-189) of LRR4 was notably moved outward (up to 6.9 Å for the Cα of C186) compared to the apo-ASK1–MAX2 structure or to the crystal structure of the ASK1–*Os*D3–*At*D14 (**Fig. 5F**), suggesting this change is linked to the presence of SMAX1.

Thus, in full-length SMAX1, the N domain stabilises the D2 domain association with MAX2 and HTL7 through an additional interaction. A protein BLAST search for *At*SMAX1 homologues in the *S. hermonthica* proteome retrieved three orthologs that we refer to as *Sh*SMXL1 (UniprotKB A0A9N7R3H5, 47% identical), *Sh*SMXL2 (UniprotKB A0A9N7NT54; 39% identical), and *Sh*SMXL3 (UniprotKB A0A9N7RQY7; 56% identical). The conservation of the N domain was stronger between SMAX1 and *Sh*SMXL1, 2, and 3 (91%, 71%, and 87% sequence identity, respectively), than between SMAX1 and *At*SMXL6 (47%), *At*SMXL7 (47%), and *Os*D53 (45%), suggesting a level of selectivity (**Supplementary Fig. 11A-C**). Structural modelling of the MAX2 complexes suggests that the N domain sequences of *At*SMXL6 and *At*SMXL7 are compatible with a MAX2 interaction, but may lead to different affinities and dynamics (**Supplementary Fig. 12A**). Indeed, previously reported HDX results indicated that *Os*D53_N_ associates with *Os*D3^53^, supporting that this N interaction with D53/MAX2 is conserved in orthologous pathways in related species.

### SMAX1 DYNAMICALLY STABILISES SCF^MAX2^ FOR UBIQUITINATION

We observed a hinge movement in the positioning of ASK1 relative to MAX2 among the cryo-EM classes (**Fig. 6A**). In the class only showing density for ASK1 and MAX2, the tip of an extended helix-turn-helix element of ASK1 (residues 74–81) contacts the MAX2 LRR11 and LRR12 opposite of its root interaction with the F-box domain (residues 35–77) and first LRR (residues 78–95) of MAX2 (**Supplementary Fig. 8D**). In this conformation (which we call the “resting state” of ASK1), the C-terminal helix element (CTH) of MAX2 (residues 704–725) was not visible, suggesting it was flexibly disengaged from the LRR domain (**Supplementary Fig. 8E**). Conversely, in the complexes with SMAX1 and HTL7, ASK1 is pulled backwards, losing the LRRs contact (the “engaged state” of ASK1). Additionally, the cryo-EM density of the ASK1 helix-turn-helix tip loses definition, indicative of increased dynamics (**Fig. 6A**, right). Concomitantly, the CTH of MAX2 becomes visible (in its helical, “engaged” state^52^) between ASK1, HTL7 and SMAX1 (**Fig. 6B)**. In the crystal structure of ASK1–*Os*D3–*At*D14 (PDB 5HZG), ASK1 is also retracted, and the CTH is engaged, suggesting that this conformation is caused by the presence of the SL receptor (D14/HTL) **(Supplementary Fig. 8A**). However, a 3D variability analysis of the cryo-EM particle stack for ASK1–MAX2–HTL–SMAX1 revealed that ASK1 engagement correlated with the presence of density for SMAX1_N_, suggesting SMAX1_N_ helps stabilise the engaged ASK1 position (**Fig. 6A, Supplementary Movie 1, Movie 2**). Concomitantly with ASK1 engagement, the density for the CTH becomes better defined and extended toward ASK1 (**Fig. 6B**). In the cryo-EM models based on Class 1 (with maximal SMAX1_N_ density), arginine residues of SMAX1_N_ are positioned to engage ionic interactions with ASK1 glutamic acids, as described above (**Fig. 5C**). Comparing apo-ASK1– MAX2 with Class 1 complex, the presence of SMAX1_N_ correlates with a 1.5 – 2 Å movement of the LRR1 and LRR2 helices toward HTL7 (**Supplementary Fig. 8F**). Both features support contacts between the MAX2 CTH and the ASK1 C-terminal helix (residues 144–160) in their engaged positions. Thus, our data support that the SMAX1 N domain stabilises the complex in the “ASK1/CTH-engaged” conformation.

**Fig. 6:**
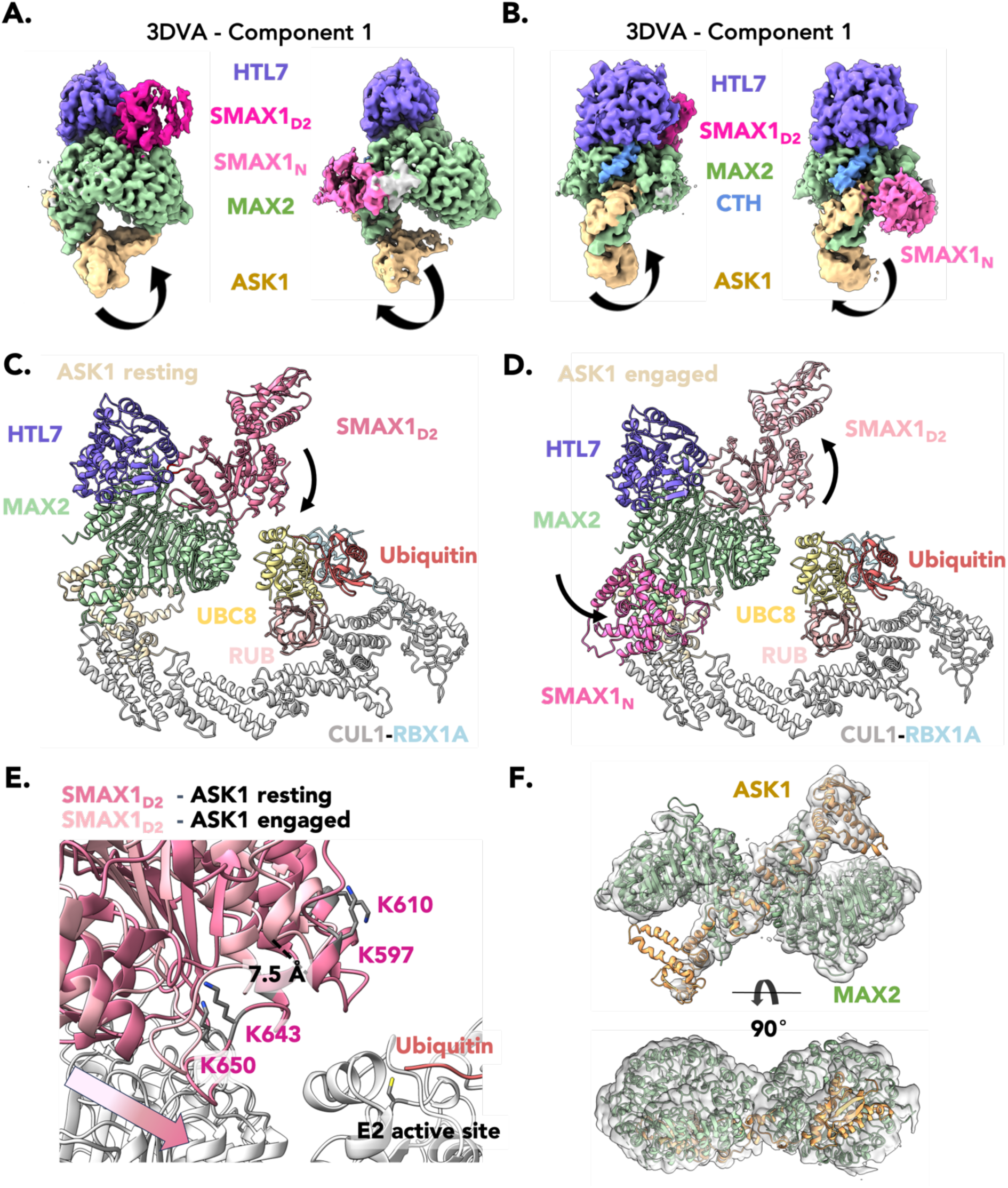
SMAX1 stabilises ASK1–MAX2–HTL7 for ubiquitination. **A.** Cryo-EM density reconstructed from the tail clusters of one variability eigenvector from 3DVA. Proteins and domains are coloured as in Fig 1. Arrows depict ASK1 motion, and correspond to **Supplementary Movie 1**. **B.** Same as in panel A, but rotated 90°. The MAX2 CTH is coloured blue. **C-D.** Model of a complete *A. thaliana* SCF–HTL7 ubiquitination complex in the resting (panel **C**) and engaged (panel **D**) ASK1 state. The ASK1–MAX2–HTL7–SMAX1 is coloured as in Fig 1. CUL1 is coloured white, its Nedd/RUB modification is coloured pale pink. RBX1A is coloured light blue. UBC8 is coloured bright yellow. Ubiquitin is coloured orange. Arrows show the relative motion between resting and engaged states. **E.** The transposition of SMAX1_D2_ in the resting (dark pink) and engaged (dark pink) ASK1 conformation and the proximity to the E2 active site and ubiquitin. The conserved lysine residues of SMAX1_D2_ are coloured grey, shown as sticks, and labelled. The distance between resting and engaged states are shown as dotted lines and labelled. The gradient arrow indicates the relative motion between the two states. The E2 active site cysteine is coloured grey and shown as sticks. **F.** Model of the 2:2 ASK1–MAX2 dimer fit in cryo-EM density (**EMD-62414**). Coloured as in Fig 1.

We next wanted to assess the impact of ASK1 movements on the positioning of the SMAX1 substrate within the full E2–SCF^MAX2^ E3–receptor–substrate complex. Using template-based modelling on PDB 8RX0^67^ (s. **Methods**), we produced the Arabidopsis complex formed by UBC8 (E2), RBX1A, CUL1, and ASK1. We then superimposed the experimental ASK1–MAX2–HTL7– SMAX1 complex onto the SCF scaffold in both the resting and engaged ASK1 states (**Fig. 6C,D**). In this context, the SMAX1 D2a subdomain is closest to Ub-conjugated E2, suggesting that ubiquitination occurs preferentially on this subdomain. Several lysines cluster in the D2a region facing the E2 ligase (K597, K610, K643 and K650), providing likely candidates for ubiquitination **(Fig. 6D)**. Ubiquitinable lysines are placed in the corresponding region in the D2 domains of other SMXLs, suggesting conservation of the mechanism (**Supplementary Fig. 12C,D**). The hinging motion of ASK1 between resting and engaged states is amplified through the CUL1 lever arm, displacing the SMAX1 D2 domain by up to 7 Å relative to the E2. The engaged state places the D2 domain further from the Ub-loaded E2, which is counterintuitive as ubiquitination is driven by proximity. However, the transient binding of the SMAX1 N domain and associated conformational changes in ASK1 would result in tapping motions and positional variations that may be required for polyubiquitination to occur.

### ASK1–MAX2 DIMERS

Although we did not observe dimerisation on SEC, our cryo-EM dataset revealed a dimeric ASK1– MAX2 class, representing a third of all particles (**Fig. 1C; Supplementary Fig. 2**). Native PAGE gels showed a weak band corresponding to a dimer, alongside a strong monomer band (**Supplementary Fig. 4A**). Mass photometry revealed a mixed population of monomers and dimers *in vitro* (**Supplementary Fig. 4B**). Despite achieving a nominal maximum resolution of 3.3 Å in 3D reconstructions, detailed structural modelling was hindered by strong anisotropy in the maps due to preferred particle orientation (**Supplementary Fig. 4C-E**). AI-based correction for anisotropy without misalignment correction (spIsoNet) led to minor improvements in map interpretability. An ASK1–MAX2 dimer model generated by AlphaFold fitted the map well, and was adjusted through rigid-body fitting. The resulting model suggested that dimerization arises from an asymmetric association between the hydrophobic N domain–binding patches on LRR1 and LRR2 of MAX2, with additional asymmetric contacts between the ASK1 C-terminal helix and the groove formed between the first and last LRR of MAX2 (**Fig 6E, Supplementary Fig. 4F**). We did not observe density for the MAX2 CTH in the dimer, suggesting it is dislodged as in the monomeric apo-ASK1–MAX2 complexes. The overlap between the dimer interface and the N domain binding site suggests competition between dimerisation and SMAX1 binding, potentially reducing the number of SMAX1-containing ASK1–MAX2(–HTL7) particles *in vitro*. The biological relevance of ASK1–MAX2 dimers remains to be determined.

## DISCUSSION

The mechanism by which SL binding and hydrolysis by their D14/HTL receptors leads to the ubiquitination of transcriptional repressors by the SCF^MAX2^ E3 ligase has remained elusive due to the transient and dynamic nature of the interactions involved. By identifying a sufficiently stable inter-species complex and preparing cryo-grids within the SCF^MAX2^ signalling time window, we obtained cryo-EM structures that reveal the conformational and compositional landscape of the SL signalling complex.

Our findings show that SMAX1 uses its N and D2 domains to engage in a dynamic bidentate association with SCF^MAX2^–HTL7. In its most fully engaged conformation, SMAX1_D2_, HTL7 and MAX2 mutually stabilise each other both directly and allosterically (by incurring the entropic penalty of MAX2 LRR17 stabilisation) on one side of the F-box LRR. Meanwhile, the SMAX1_N_ domain (and possibly an additional unidentified SMAX1 region), MAX2, and ASK1 interact on another MAX2 surface. Although separated by more than 10 Å, both regions appear mechanistically connected through the MAX2 CTH in its “engaged” helical form. Our structural and biochemical analyses show that the individual association of SMAX1_N_ and SMAX1_D2_ with MAX2–HTL7 are weak, leading to fluctuations around the bound position and temporary detachment of one or the other. The lack of structural coupling between the N and D2 domains allows the bidentate interaction to avoid excessive affinity gains or structural rigidity through avidity, while achieving overall higher affinity and specificity due to increased local concentrations of one domain when the other is bound.

We show that the presence of density for SMAX1_N_ correlates with the retraction of ASK1 from its “resting” position on MAX2. Within the fully assembled E2–E3–substrate complex, the associated ASK1 movements are amplified by the long cullin lever arm. N domain binding dynamics would result in a tapping encounter between the ubiquitin-conjugated E2 and the D2 domain. We hypothesise that these structural dynamics are essential for efficient substrate polyubiquitination. Our analysis provides a mechanistic explanation for previous indirect observations that dynamics in the SCF^MAX2^–receptor–substrate complex are crucial for substrate polyubiquitination and release^49,51,52,68^. Additionally, our findings suggest that the SMAX1 N domain uses its MAX2-binding surface to interact with TCP-family transcription factors (**Supplementary Fig. 10C**)^64^. Hence, the bidentate association may help the SCF^MAX2^–receptor complex to disengage SMAX1 from TCP family protein.

The involvement of the D2 domain in the SCF^MAX2^–SL signalling complex has been strongly suggested by various lines of evidence^52,55,61^. Our results provide a mechanistic explanation for these observations. We demonstrate that SMAX1_D2_ has been repurposed from an ATPase domain into a binding module through the loss of its nucleotide binding site and the use of its pore loop for allosteric stabilisation of the association between the F-box and the SL-bound receptor. Thereby, our structures revealed the role of the conserved RGKT motif in stabilising the complex both directly and allosterically.

The role and timing of SL cleavage during SCF^MAX2^–D14/HTL signalling and the nature of the SL cleavage product associated with the SCF^MAX2^–D14/HTL–substrate complex has remained controversial. Considering previous evidence that SL binding destabilises the receptors^29,30^, our structural and HDX results indeed show that the receptor lid only stably closes upon binding to the F-box. In this state, SL cleavage must be completed due to the space constraints. However, our results are compatible with an initial encounter complex between the dislodged MAX2 CTH and a non-collapsed SL-bound receptor^52^, which then transitions to the fully assembled SCF^MAX2^– receptor–substrate complex where CTH is engaged with MAX2. Our results also do not rule out a biological role of lid-open D14/HTL bound to an intact SL^29^, but it would need to occur prior to fully engaging the MAX2 LRR surface.

Features of the electron density in the crystal structure of the ASK1–*Os*D3–*At*D14 complex (PDB 5HZG) led the authors to conclude that the SL cleavage product is the open D-ring covalently bound to both the catalytic Ser and His of D14^28^. However, subsequent reanalysis of these data by other researchers refuted this claim, suggesting that the observed electron density rather corresponds to either an iodide or a closed D-ring bound to the catalytic His^47,48^. Our findings strongly support that the closed D-ring, covalently attached to the catalytic His246, is the receptor-associated SL cleavage product during the signalling stage. This visual conclusion is in line with previous biochemical evidence^28,56,59^. Our cryo-EM density fully resolves the catalytic centre of the HTL7 receptor, including the D-loop, which was not visible in the *At*ASK1–*Os*D3– *At*D14 crystal structure^28^. The resulting structure reveals that the catalytic triad of HTL7 is sufficiently distorted in the complex with MAX2 and SMAX1_N_ to halt its catalytic activity. Thus, we concluded that SMAX1 stabilises the receptor in a catalytically inactive conformation.

Although D14 and HTLs (and the homologous KAI2) engage SCF^MAX2^, they trigger different biological outcomes, with only HTL/KAI2 being associated with germination^44,69,70^. Previous work evidenced a level of selectivity of the SCF^MAX2^–D14 for the substrates SMXL6, SMXL7, and SMXL8, whereas SCF^MAX2^–HTL/KAI2 preferentially targets SMAX1 and SMXL2^32,61^. Despite significant efforts from several groups, the determinants for specificity remained ill-defined and controversial, with N, D1, M, D2 domains and combinations thereof, proposed as key elements^35,52,61^. Moreover, in many experiments, the selection of one substrate over another appears preferential rather than stringent. Indeed, we observed that GR24-supplemented GST-HTL7 co-precipitated ASK1–MAX2 not only in the presence of SMAX1 but also in the presence of the D14-specific SMXL6 and SMXL7. According to our structural analysis, only the D2 domain of SMAX1 is in direct contact with the HTL7 receptor. These contacts are relatively small (430 Å^2^), but match better within the SMAX1–HTL7 and SMXL7–D14 pairs than across pairs, according to homology modelling (**Fig. 7A**). Although our structures were prepared with full-length SMAX1, we did not see other density in contact with HTL7 to indicate additional determinants of receptor specificity. The only non-attributed density is adjacent to the N domain, away from the D14/HTL7 binding site on MAX2. The N domains of SMAX1-LIKEs have been shown to be involved in developmental control in *A. thaliana*^64^. Our analyses support that the N domains of *At*SMXL1, 6 and 7 can associate with MAX2 in the same way, but with altered affinities and dynamics. Therefore, we conclude that it is the sum of individual affinities that determine the stability and dynamics of the SCF^MAX2^–receptor–substrate complex, which in turn translate into substrate ubiquitination efficiency and thus substrate selectivity and biological function. While the receptor–substrate match is a significant component of this ensemble, it is not sufficient on its own.

**Fig. 7:**
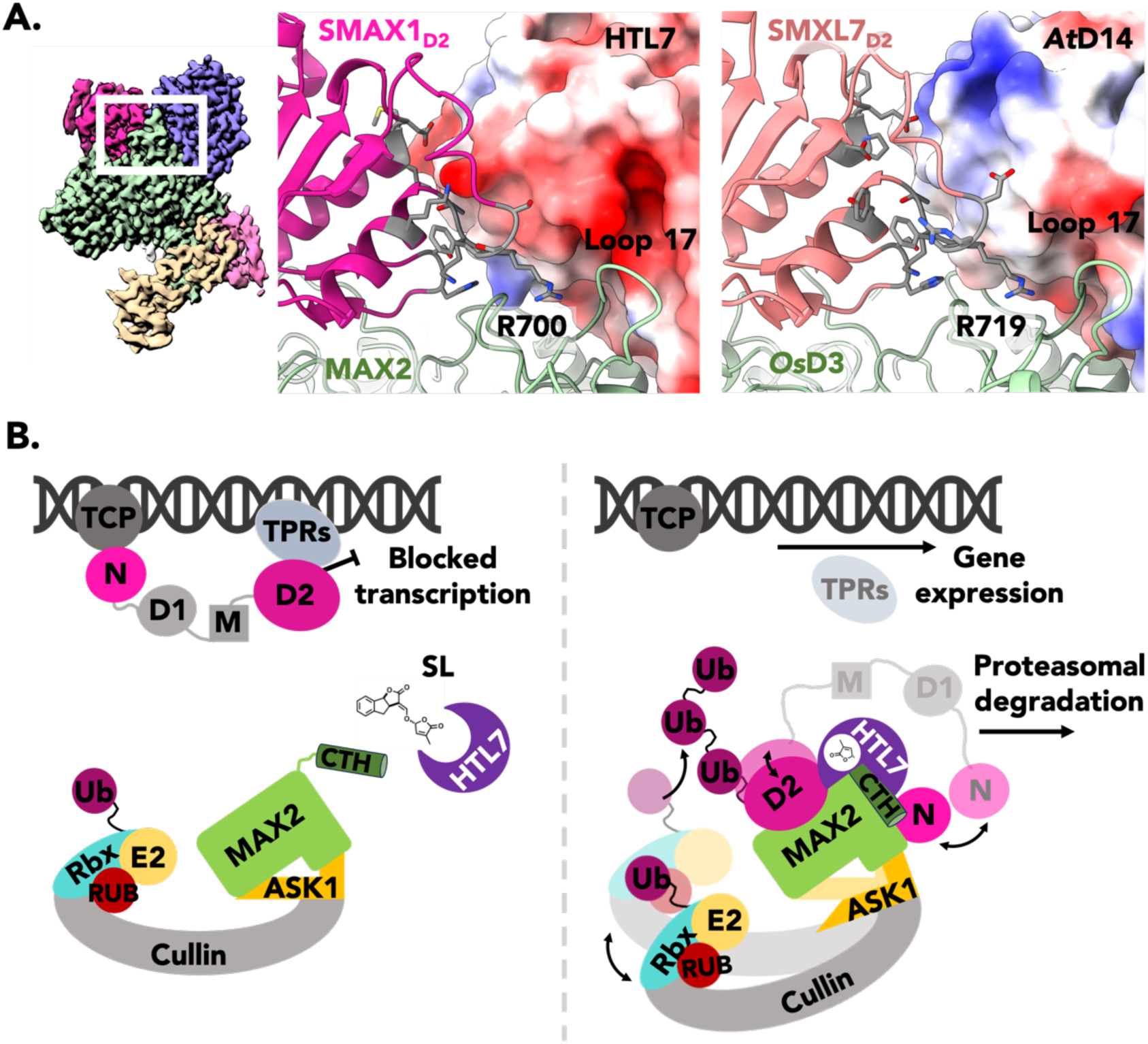
Receptor contributions to specificity and schematic overview. **A.** Reference view (left) of ASK1–MAX2–HTL7–SMAX1 (**EMD-62417**) with a white box outlining the focal area. Interface of MAX2–HTL7–SMAX1_D2_ (middle, **9KLD**) and *Os*D3–*At*D14–SMXL7_D2_ (right, PDB 5HZG). The SMAX1_D2_/SMXL7_D2_ (dark pink/light pink) RGKT motif and interface residues with HTL7/*At*D14 are shown as sticks and coloured grey. HTL7/*At*D14 are shown as a surface and coloured by electrostatic potential from red (negatively charged) to blue (positively charged). MAX2/*Os*D3 are coloured green. **B.** Updated schematic of the strigolactone signalling mechanism. In the pre-signalling state (left), SMAX1 is bound to TCP and TPRs, blocking transcription. The MAX2 CTH is dislodged, as seen in apo-ASK1–MAX2 (**PDB 9KLL**). Strigolactone perception and hydrolysis enables HTL7 to close and bind to MAX2, trapping the D-ring in the active site. This HTL7 conformation creates the interface for SMAX1_D2_ to mutually stabilise the tripartite MAX2–HTL7– D2 complex. Simultaneously SMAX1_N_ engages another site on the MAX2 LRR, close to the MAX2 CTH and ASK1, promoting the “engaged” position of ASK1. The transient interactions of SMAX1_N_ lead to a tapping motion of the E2 towards SMAX1_D2_, while the interaction dynamics of SMAX1_D2_ with MAX2 lead to positional variations towards the E2. Together, these oscillating interactions flexibly position SMAX1_D2_ for polyubiquitination, leading to proteasomal degradation of SMAX1, thereby enabling transcription to occur.

In addition to the controlled co-expression of the dozens of proteins involved in substrate ubiquitination and degradation^71^, the SL perception machinery may also be sensitive to additional regulatory layers, such as small molecules (e.g., citrate or inositol phosphates)^19,49,51^ or post-translational modifications^72^, which could enhance environmental sensing. The complexity of the SCF^MAX2^-based SL perception, which, remarkably, occurs in Striga only after decade-long seed dormancy and conditioning, may reflect the level of signal integration required for Striga to commit to germination only in the presence of a host and under favourable conditions.

In summary, our work fills an important knowledge gap by capturing the long-elusive SCF^MAX2^–receptor–substrate complex that mediates SL perception. Our results help rationalise and resolve previous observations pertaining to the mechanistics of SL perception, and may inspire chemical or genetic interventions to control Striga germination. Additionally, these findings highlight the importance of finely tuned dynamics in E3-based hormone perception (**Fig. 7B**). Such dynamics are likely underappreciated in other systems, due to the challenge of visualising them through structural analyses that rely on averaging many molecules (e.g., X-ray crystallography and cryo-EM). Thus, our work significantly advances our understanding of how E3 ligases are used in plants to translate hormone perception into genetic adaptations.

## MATERIALS AND METHODS

### Plasmid construction

The Multibac™ expression system from Geneva Biotech was used to co-express in insect cells each of the full-length proteins *Os*D3, *At*MAX2, and *Sh*MAX2 with *At*ASK1. The genes encoding *Os*D3 (UniprotKB Q5VMP0), *At*MAX2 (UniprotKB Q9SIM9), *Sh*MAX2 (UniprotKB T1RVG4) and *At*ASK1 (UniprotKB Q39255) were ordered from Twist Bioscience, with an N-terminal cleavable tandem 9xHis tag and Strep-tag for the *At*MAX2 and *Sh*MAX2 inserts. *Os*D3 insert was amplified with an N-terminal cleavable Strep-tag and a C-terminal 6xHis tag and was cloned in the pIDK donor vector. The *At*MAX2 and *Sh*MAX2 inserts were cloned in the pACEBac1 acceptor vector and the untagged *At*ASK1 insert was cloned in both pIDS and pACEBac1 vectors. The single transfer vectors with different MAX2 and *At*ASK1 inserts were generated using Cre recombinase according to the MultiBac™ expression system user manual, and the recombinant transfer vectors were transformed into DH10EMBacY™ cells, with a constitutively expressing YFP expression cassette. The resulting bacmid DNA was isolated by isopropanol precipitation.

Full-length *At*SMAX1 gene (UniprotKB Q9FHH2) was ordered from Twist Bioscience and cloned into the pQlinkH vector (with an N-terminal 7xHis tag) by restriction cloning between the BamHI and NotI restriction sites. An additional C-terminal Strep-tag was added by overhang PCR amplification with phosphorylated primers, followed by ligation. The corresponding *At*SMAX1 truncations and mutant constructs were generated similarly, by PCR amplification with phosphorylated primers. *At*SMXL6 (UniprotKB Q9LML2) and *At*SMXL7 (UniprotKB O80875) were ordered from Twist Bioscience with an N-terminal tandem Strep and a C-terminal 7xHis tag in pJex vectors. *Sh*HTL7 gene (UniprotKB A0A0M3PNA2) cDNA^60^ was cloned into pQlinkH and pGEX6P-1 (with an N-terminal GST tag) vectors by restriction cloning between the BamHI and NotI restriction sites.

### Protein preparation

For insect cell expression, Sf9 cells were cultured in ESF 921 medium (Expression Systems) at 27°C. The bacmid DNA was transfected into Sf9 cells in adherent culture using FuGENE® HD (Promega), and the YFP fluorescence was monitored for baculovirus production. The baculovirus was amplified twice in suspension culture to obtain a higher viral titer. For the expression of the MAX2–ASK1 proteins, the amplified virus was used to infect Sf9 cell cultures at a density of 2 × 10^6^ cells/mL. The cells were harvested at 70-80% cell death post-infection within 66-72 h by centrifugation and the pellet was gently washed with PBS or purification buffer and was stored at −80 °C if purification was not done immediately after.

The SMAX1 plasmid was transformed into *E. coli* BL21 (DE3) cells grown on agar plates with ampicillin selection. A single colony was used to grow an overnight starter culture in 2YT medium. For expression, flasks containing 1 L culture media were inoculated with the starter culture and the cells were grown at 37 °C and 200 rpm to reach the OD600 of 0.6, when protein expression was induced by adding isopropyl β-D-1-thiogalactopyranoside (IPTG) to 0.2 mM at 16°C for 12-16 h. A similar protocol was used for expressing *At*SMXL6, *At*SMXL7 and *Sh*HTL7 from BL21 (DE3) cells. LB media was used for ShHTL7 expression.

For all MAX2 and *At*SMAX1 proteins purification was done on ÄKTA pure™ systems, first by affinity purification on HisTrap™ HP 5 mL (Cytiva) column, followed by StrepTrap™ XT 5 mL (Cytiva) column and then by size-exclusion chromatography (SEC) on HiLoad 16/600 Superdex 200 pg (Cytiva). *At*ASK1 co-eluted with all MAX2 proteins after all the purification steps. Cell pellets were first homogenized in binding buffer (50 mM Tris-HCl pH 7.5, 500 mM NaCl, 1 mM TCEP, 20 mM imidazole, 4% (v/v) glycerol) with protease inhibitors (SIGMAFAST™ EDTA free) and benzonase (MilliporeBenzonase® Nuclease) by gentle sonication or using a manual cell homogenizer, and then lysed using a cell disruptor. The lysate was clarified by centrifugation at 50,000 g and passed through the His-affinity column equilibrated with binding buffer and the protein was eluted with 250 mM imidazole after an initial washing step with 40 mM imidazole. The protein fractions were then loaded on the Step-affinity column equilibrated with binding buffer (with adjusted imidazole composition) and eluted after washing with 50 mM D-(+)-Biotin (Thermo Scientific) after 1 h incubation on bead. The protein fractions were injected into the SEC column equilibrated with 20 mM HEPES pH 7.5, 200 mM NaCl, 1 mM TCEP and 4% (v/v) glycerol (for *At*SMAX1 300 mM NaCl was used) and the pure fractions were concentrated and snap-frozen for storage at −80 °C. An SDS-PAGE gel was run after each purification step. All purification steps were performed at 4 °C and as quickly as possible, particularly for *At*SMAX1 which showed signs of degradation.

His-tagged and GST-tagged *Sh*HTL7 were purified by gravity flow chromatography in glass Econo-Column® from Bio-Rad. Cell pellets expressing His-tagged *Sh*HTL7 were lysed by sonication in binding buffer containing 50 mM Tris-HCl pH 8.0, 200 mM NaCl, 1 mM DTT, 20 mM imidazole with 0.1% (v/v) Tween-20 added, and the clarified lysate was loaded onto Ni-NTA Agarose resin (QIAGEN). The resin was washed with binding buffer containing 40 mM imidazole and eluted with binding buffer containing 250 mM imidazole. The eluted protein fractions were further purified on the SEC column equilibrated with 20 mM HEPES pH 7.5, 150 mM NaCl, and 3 mM DTT. Protein purity was evaluated using SDS-PAGE. The purified proteins were concentrated and stored at −80°C. The GST-tagged *Sh*HTL7 was purified similarly using 50 mM Tris-HCl pH 8.0 200 mM NaCl, 1 mM DTT with 0.1% (v/v) Tween-20 added for lysis, and loaded on Glutathione Sepharose 4B (Cytiva) resin. After washing with the binding buffer, the protein was eluted with 20 mM reduced glutathione (VWR) in the binding buffer, followed by SEC on HiLoad 16/600 Superdex 75 pg (Cytiva). The pure fractions were concentrated and snap-frozen for storage at −80 °C.

### Size-exclusion chromatography (SEC) assay

Purified *Sh*MAX2–*At*ASK1 (17 μM), *Sh*HTL7 (25 μM) and *At*SMAX1 (20 μM) in 150 μL reaction volume were incubated at 4 °C for 30 min with 200 μM GR24 or an equal amount of DMSO as the solvent control, in buffer containing 20 mM HEPES pH 7.5, 200 mM NaCl, 1 mM TCEP. The reactions were injected into a Superdex 200 10/300 column (Cytiva) for analysis at a flow rate of 0.5 mL/min. The fractions were analysed by SDS-PAGE and visualised by Coomassie staining. For testing the interaction of other MAX2–HTL(–SMXL) components, the proteins were mixed in a 1:1.5:1.2 molar ratio to a final 150 μL volume with roughly 0.25 mg of the MAX2 protein, and corresponding amounts of the binding partners, following the same procedure as above.

### Pull-down assay

For testing the *Sh*HTL7 interaction with *Sh*MAX2–*At*ASK1 and *At*SMAX1 (full length, D2 domain and corresponding mutants) or *At*SMXL6/*At*SMXL7 using GST-*Sh*HTL7 as bait, reaction mixtures of 100 µL in 20 mM HEPES pH 7.5, 200 mM NaCl, 0.5 mM TCEP buffer were prepared for a final concentration of roughly 3.5 µM GST-*Sh*HTL7 and 5 µM for each of the tested binding partners. Purified GST-*Sh*HTL7 was first bound to beads (Glutathione Sepharose 4B resins) and the beads were aliquoted to each reaction mixture. The reactions were incubated with 100 µM GR24, MP3 or their solvent DMSO as the control for 30 minutes on a rotator at 4 °C and protected from light. The beads were then rinsed with 150 µL buffer and washed three more times with 5 min incubation on the rotator, before elution with 40 µL buffer with 20 mM glutathione by incubation for 20 min. The washes were monitored by Bradford assay (Bio-Rad) and continued until no protein was visibly detected in the flow-through. The eluted fractions were analysed by SDS-PAGE. A similar procedure was used for testing *Sh*HTL7 interaction with *Os*D3–*At*ASK1 with GR24 and MP3. The eluted fractions were tested for *Os*D3 presence by western blot using anti-6xHis HRP-conjugated antibody (Abcam). For testing *Os*D3–*At*ASK1 interaction *Sh*HTL7 using Strep-tagged *Os*D3 as bait, 200 µL reaction mixtures (in the same buffer) of the two proteins of roughly 10 µM *Os*D3–*At*ASK1 and 17 µM *Sh*HTL7 were prepared and incubated for 30 min with GR24 and MP3, and then loaded to Strep-Tactin® XT beads and incubated for another 30 min. Washes were performed as above and the proteins were eluted in buffer with 50 mM biotin, and the fractions were analysed by SDS-PAGE.

### Cryo-EM specimen preparation

Cryo-EM grids (Quantifoil R1.2/1.3, 400 mesh Au or UltrAuFoil, R1.2/1.3, 300 mesh) were washed with acetone for 30 s, isopropyl alcohol for 10 s and left to dry. Washed grids were then glow discharged in air using either a Quorum GloQube (60 s at 35 mA current) or a Pelco easiGlow (30 s at 30 mA current). 3 µL of the sample (*At*ASK1–*Os*D3–*Sh*HTL7 at 0.5 mg/mL or *At*ASK1–*Sh*MAX2– *Sh*HTL7–*At*SMAX1 at 0.35 mg/mL) was applied to freshly glow-discharged grids, blotted for 2-3 s and plunge-frozen using a TFS Vitrobot Mark IV at 4 °C and 100% humidity. Samples with detergent were mixed with detergent (final 0.005% (v/v) Tween-20) just before vitrification.

### Cryo-EM data collection

Cryo-EM data were recorded on a TFS Krios G4 operated at 300 kV, equipped with a Selectris X energy filter and Falcon 4i direct detector. For the *At*ASK1–*Sh*MAX2–*Sh*HTL7–*At*SMAX1 dataset, 20,175 EER movies were collected at 165k magnification, yielding a pixel size of 0.73 Å/pixel at specimen level. A flux of 7.8 e^-^/pixel/s resulted in a total fluence of 44 e^-^/Å^2^ on the specimen, over a 3 s exposure time. Data were collected using aberration-free image shift (AFIS) of +/-12 µm in EPU (v3.5) with a nominal defocus range −2.7 µm to −1.5 µm in 0.3 µm intervals and an energy filter slit width of 10 eV, with automatic re-centring of the zero-loss peak every 2 hours.

For the *At*ASK1–*Os*D3–*Sh*HTL7 untilted data set of 6,741 movies were recorded similarly, at a flux of 4.76 e^-^/pixel/s resulting in total fluence on the specimen of 40 e^-^/Å^2^ over a 4.5 s exposure time. Similar collection parameters were used for the tilted dataset (7,372 movies), except for a reduction of AFIS to +/-6 µm, a single nominal defocus value of −2 µm, and a +30° stage tilt.

### Cryo-EM image processing

For the *At*ASK1–*Sh*MAX2–*Sh*HTL7–*At*SMAX1 dataset, an overview of the data processing can be found in **Supplementary** Fig 2. Initial processing was performed in RELION (version 5.0-b commit-90d239)^73^. Movies were motion-corrected using RelionCorr with an EER fractionation of 21 and electron exposure of 1.02 e^-^/Å^2^ per fraction. CTF estimation was carried out using CTFFind4^74^ and images were curated based on resolution (<6 Å), defocus (−1.0 to −3.0 µm), and relative figure of merit resulting in 15,184 accepted images. Particles were picked using the default model of Topaz^75^ on a subset of 1k images to generate references for template-based picking. After picking on the entire data set by template - based picking and doing several rounds of 2D classification-based cleaning on particles extracted with 360 pix and 4x binning we obtained a particle stack of 1.8 m particles. The cleaned particle stack was re-extracted with 2x binning and imported into CryoSPARC (version 4.4)^76^. Multi-class Ab-Initio Reconstruction followed by Homogeneous Refinement and 3D Classification jobs were used to generate high-quality reference volumes for each compositional assembly. Additionally, 4 decoy classes of pure noise were generated by sampling a very small number of particles using Ab-Initio Reconstruction. Concomitantly, the images from RelionCorr were imported into CryoSPARC and curated similarly. The images were over-picked using blob picking (diameter 80 Å to 170 Å) to generate a particle stack of 7.4 m particles with a box of 360 pix with 2x binning.

The over-picked particle stack with high-quality references and decoy volumes was used for iterative Heterogeneous Refinement in CryoSPARC. Only particles retained in the high-quality reference classes were used as input for each sequential Heterogeneous Refinement until >90% of particles were retained in these classes. Particles corresponding to each compositional assembly were grouped and extracted without binning and Non-Uniform Refinement (C1 with global and local CTF refinements) was performed to generate the consensus refinements.

These consensus particle stacks were imported back to RELION using pyem^77^ where alignment-free 3D classification using Blush^78^ was used to separate different conformations. For selected classes, iterative CTF Refinement, Bayesian Polishing, and Refine3D using Blush and local searches only was performed to generate the final volumes. For Class 2 of the *At*ASK1–*Sh*MAX2– *Sh*HTL7 complex, an additional round of alignment-free 3D classification was performed with a large mask around *At*SMAX1_N_ to identify a subset of particles with the best *At*SMAX1_N_ density. The Dimer particle stacked comprised primarily side views, causing anisotropic reconstructions. Selecting classes with highest isotropy and attempting to rebalancing orientations were unsuccessful at alleviating the anisotropy.

To improve the interpretability of maps for the initial stages of model building, unsharpened maps were subjected to the EMReady algorithm^79^. To improve the isotropy of the Dimer map, the unsharpened map was subjected to Anisotropy Correction without Misalignment Correction using the SpIsoNet algorithm^80^. The corrected Dimer map was then further modified using EMReady to improve the interpretability.

The untilted and tilted *At*ASK1–*Os*D3–*Sh*HTL datasets were processed in RELION (version 4.0.0-commit-138b9c)^73^. Both data sets were processed independently to the stage of 3D classification before joining the curated particle stacks for further processing. An overview of the data processing can be found in **Supplementary** Fig 1. Movies were motion-corrected using RelionCorr with an EER fractionation of 33 and an electron exposure of 0.96 e^-^/Å^2^ per fraction. CTF estimation was carried out using CTFFind4^74^ and images were curated based on resolution (<6 Å) and relative figure of merit resulting in 5,949 accepted images for the untilted and 3,175 movies for the tilted datasets. The default model of Topaz^75^ was used to pick particles, which were extracted with a box size of 220 pix with 2x binning. Particles were curated through iterative 2D and 3D classifications, yielding a final stack of 227k particles. Refinement of this particle stack reached 4.4 Å, which was further improved to 4.3 Å after CTF refinement. Further polishing or CTF refinement did not improve the quality of the map.

### Cryo-EM variability analysis

3DVA^81^ on *At*ASK1–*Sh*MAX2–*Sh*HTL7–*At*SMAX1 was performed from the consensus refinement using a filter resolution of 6 Å and mask containing all domains. The heterogeneity was represented along 3 eigenvectors. 3DVA Display in Simple mode was used to generate 20 volumes along each eigenvector. Movies were then generated for each volume series in ChimeraX^82^. To discretise the heterogeneity, 3DVA Display in Cluster mode was used to generate 5 clusters. The tail clusters from the variability distribution along each eigenvector were refined using Local Refinement with the same mask. These end-states were used to interpret the extreme ranges of heterogeneity present in the particle stack. 3DVA of apo-ASK1–MAX2 did not suggest substantial heterogeneity. Due to the high anisotropy of the Dimer, 3DVA was not performed.

### Model generation, fitting and refinement

Initial models were generated with AlphaFold^83^ and SWISS-MODEL^84^ using 5HZG as a template for the collapsed form of HTL7. Models were rigid-body docked into the cryo-EM densities using ChimeraX^82^ and flexibly fit using Isolde^85^. Iterative manual adjustments of the models were performed in Coot^86^ alongside optimisation of the geometry and fit to density through Real Space Refinement in Phenix^87^, including the ligand restraint file when necessary. The restraint file for the *Sh*HTL7 ligand (D-ring) was generated using eLBOW^88^ after using the Ligand Builder tool in Coot^86^ to generate the SMILES string.

### Model of the E2–SCF^MAX2^ E3–receptor–substrate complex

The SCF–E2–Ubiquitin scaffold was obtained by assembling Arabidopsis CUL1 and RBX1A (modelled with SWISS-MODEL^84^ on the template PDB 8RX0), with the ubiquitin chain (chain U) from PDB 8RX0, the Arabidopsis E2 UBC8 attached to an Ub-like molecule (chains A and B of PDB 4X57) which superimposed well (Cα RMSD 0.751 Å) with the E2 and NEDD8 of 8RX0 (chains N and G), and *At*ASK1 from the apo-*At*ASK1–*Sh*MAX2 model. The PDB 8OR0 (structure of a *Hs*Cullin-1 complex) superimposed onto the *Hs*Cullin-2 (chain E) of PDB 8RX0 (Cα RMSD 1.224 Å) was used as reference to position ASK1 on the modelled *At*CUL1. The experimental *At*ASK1–*Sh*MAX2– *Sh*HTL7–*At*SMAX1 complex was superimposed onto the generated SCF scaffold and the full-length AlphaFold predicted model of *At*SMAX1_D2_ was used.

### Hydrogen/deuterium exchange mass spectrometry (HDX-MS)

The individually purified *Sh*HTL7, *At*SMAX1 and *Sh*MAX2–*At*ASK1 proteins were dialyzed in 20 mM HEPES pH 7.5, 200 mM NaCl, 0.5 mM TCEP buffer and then mixed in equimolar ratios to reach final protein concentrations of 25 µM in 200 µL (see **Supplementary Dataset 1**). Prior to HDX-MS, the protein stock solution was supplemented with 0.8 µL of 50 mM GR24 dissolved in DMSO or a DMSO control yielding 200 µM GR24 final concentration. This stock solution was temperated at 1 °C for the entire HDX data acquisition time of approx. 24 h. The preparation of individual HDX reactions was aided by a two-arm robotic autosampler (LEAP Technologies) as described previously^89^. In brief, 7.5 *μ*L of sample stock solution were pre-dispensed in a plate temperated at 25 °C, then 67.5 *μ*L of HDX buffer (20 mM HEPES-Na pH 7.5, 200 mM NaCl, 0.5 mM TCEP) prepared with 99.9% D_2_O added and incubated for 10, 100, 1,000 or 10,000 s at 25 °C. 55 µL of the HDX reaction were then withdrawn and add to 55 µL of quench buffer (400 mM KH_2_PO_4_/H_3_PO_4_, pH 2.2, 2 M guanidine-HCl) pre-dispensed in another plate and kept at 1 °C. After mixing, 95 µL of the quenched reaction were injected into an ACQUITY UPLC M-Class System with HDX Technology (Waters)^90^. Non-deuterated HDX samples were prepared similarly by 10-fold dilution of sample stock solutions with HDX buffer prepared with H_2_O (incubation for approximately 10 s at 25 °C). The protein samples were digested online with porcine pepsin and separated by reversed-phase HPLC followed by mass spectrometric analysis of three technical replicates (individual HDX reactions) per HDX time-point as described^89^.

Peptides were identified from the non-deuterated samples with the ProteinLynx Global Server 3.0.1 (Waters) with a database containing the amino acid sequences of *Sh*HTL7, *At*SMAX1, *Sh*MAX2, *At*ASK1, porcine pepsin, and their reversed sequences. For quantification of deuterium incorporation with DynamX 3.0 (Waters), peptides had to fulfil the following criteria: minimum intensity of 10,000 counts; length of 5-40 amino acids; minimum number of products of two; maximum mass error of 25 ppm; retention time tolerance of 0.5 min. After automated data processing by DynamX 3.0, the mass spectra were manually inspected. The observable maximal deuterium uptake of a peptide (**Supplementary Dataset 1**) was calculated by the number of amino acid residues minus one (for the N-terminal residue) minus the number of proline residues. For rendering of the residue-specific HDX and differences thereof from overlapping peptides, the shortest peptide covering a residue was employed. Where multiple peptides were of the shortest length, the peptide with the residue closest to the peptide’s C-terminus was utilised. Peptide and residue-level HDX values are provided as **Supplementary Dataset 1**.

## ACKNOWLEDGMENTS

We thank L. Zhao for help with cryo-EM data recording. We thank S. Al-Babili for valuable discussions and reagents. For computer time, this research used the resources of the KAUST Supercomputing Laboratory, and experimental research was supported by the Bioscience Core Lab, ACL Proteomics Lab and the Imaging and Characterisation Core Lab at King Abdullah University of Science & Technology (KAUST) in Thuwal, Saudi Arabia.

## AUTHORS CONTRIBUTIONS

Study design: A.I.V, U.F.S.H, S.T.A. Protein production and biochemical analysis: A.I.V. HDX data collection: W.S. HDX analysis: A.I.V., W.S., S.T.A. Cryo-EM sample preparation and data collection: A.I.V., C.S. Biochemical and structural data analysis: A.I.V., B.H., C.S., S.T.A. Manuscript writing: A.I.V., B.H., S.T.A. All authors read the manuscript and provided comments.

