## Supplementary Figures for "Mechanism of cooperative strigolactone perception by the MAX2 ubiquitin ligase–receptor–substrate complex"

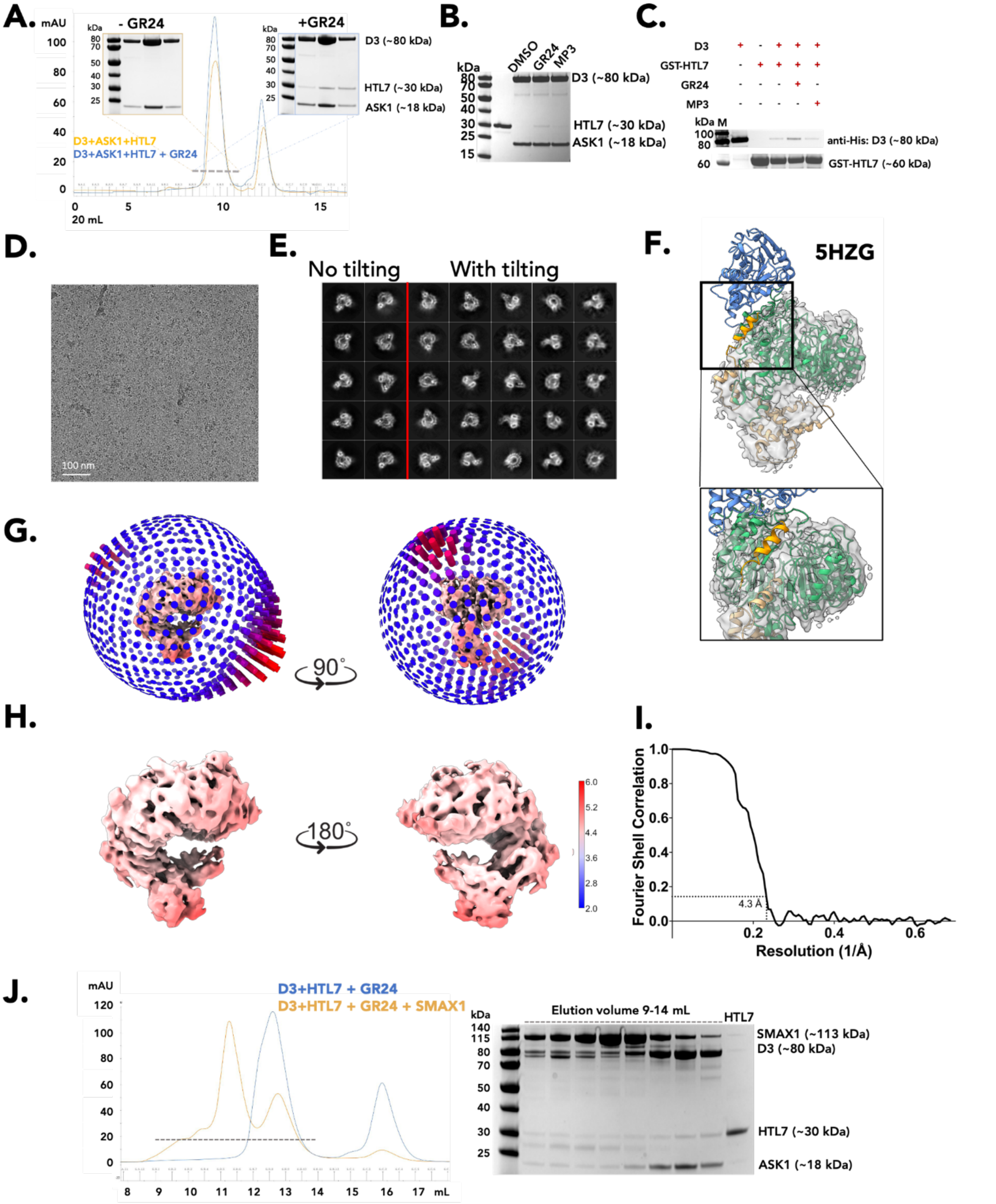


**Supplementary Fig. 1: Biochemical and structural characterisation of ASK1–*Os*D3–HTL7. A.** SEC profiles and corresponding Coomassie stained SDS-PAGE of ASK1–*Os*D3–HTL7 with (blue; right) and without (orange; left) GR24. Solid lines show the absorbance at 280 nm. Grey dashed lines highlight the fractions analysed by SDS-PAGE. The corresponding SDS-PAGE gel for each SEC profile is outlined with the same colour. Approximate molecular weights are labelled. Uncropped gel (blue outline) shown in Supplementary Fig. 13. **B.** Coomassie stained SDS-PAGE of pull-down assays using Strep–*Os*D3 as a bait and HTL7 as the target in the presence of DMSO, GR24, or MP3. Lane 1 shows the marker and Lane 2 corresponds to a control with HTL7 alone. Approximate molecular weights are labelled. **C.** Western blot analysis of GST pull-down assays using GST-HTL7 as a bait and combinations of *Os*D3, GR24, and MP3. Plus signs denote presence and minus signs absence of species in each pull-down assay. Anti-6xHis antibodies were used to detect *Os*D3. Lane below shows the Coomassie stained SDS-PAGE for the loading of the GST-HTL7 bait on the beads. "M" denotes the marker lane. Approximate molecular weights are labelled. Uncropped gel and membrane shown in Supplementary Fig. 13. **D.** Representative cryo-EM image from the ASK1–*Os*D3–HTL7 dataset. **E.** Combined 2D class averages of untilted (left) and tilted (right) ASK1–*Os*D3–HTL7 cryo-EM datasets. Red line delineates the two datasets. **F.** Cryo-EM density of ASK1–*Os*D3–HTL7 (**EMD-62415**) as a transparent grey surface with PDB 5HZG overlaid. ASK1, *Os*D3, and *At*D14 are coloured yellow, green, and blue, respectively. The CTH of *Os*D3 is coloured gold. Black box shows a rotated and magnified view of the CTH. No cryo-EM density for HTL7 or the CTH is visible **G.** Euler angle distribution of the final ASK1–*Os*D3–HTL7 reconstruction. **H.** Cryo-EM density coloured by local resolution. Higher resolutions are coloured blue and lower resolutions are coloured red. **I.** Fourier shell correlation (FSC) curves for the cryo-EM reconstructions. Black line represents the correlation between independently processed half-maps, with resolution determined at FSC = 0.143 threshold. **J.** SEC profiles (left) of ASK1–*Os*D3–HTL7 and GR24 with (orange) and without (blue) SMAX1. Solid lines show the absorbance at 280 nm. Black dashed lines highlight the fractions analysed by SDS-PAGE. SDS-PAGE (right) of ASK1–*Os*D3–HTL7 and GR24 with SMAX1. Control lane of HTL7 is also shown. Approximate molecular weights are labelled.

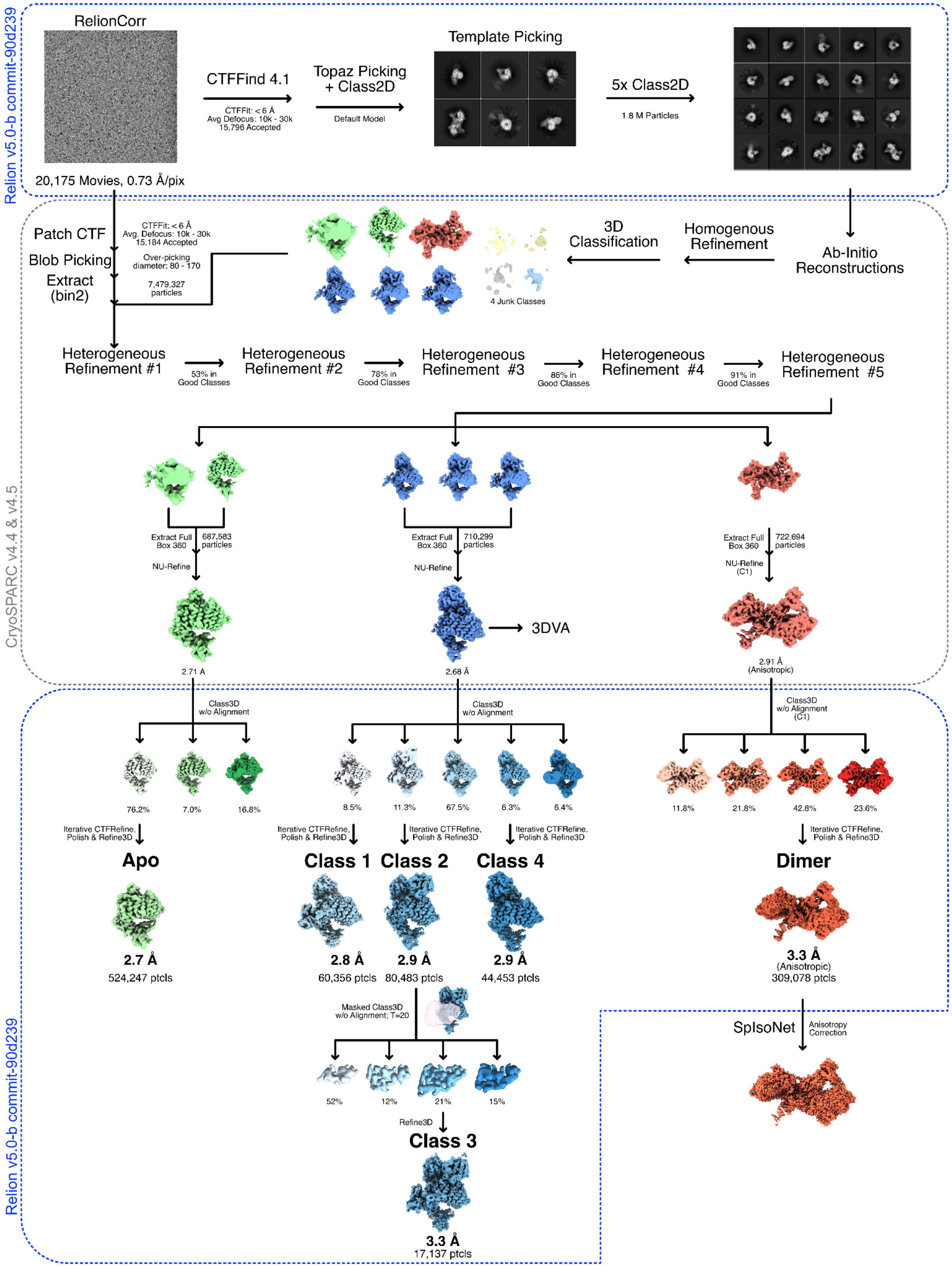


**Supplementary Fig. 2: Cryo-EM image processing diagram of ASK1–MAX2–HTL7–SMAX1.**

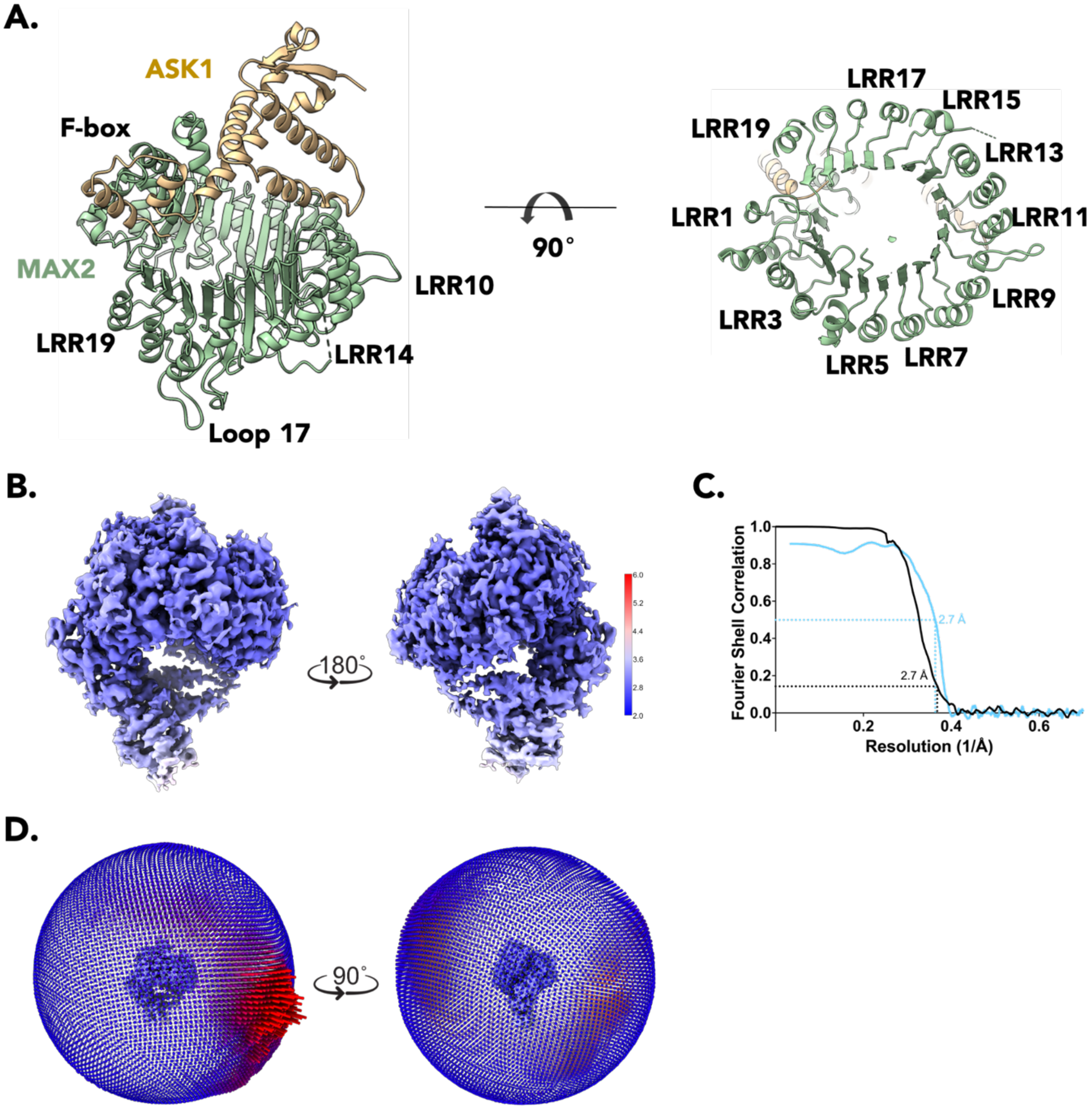


**Supplementary Fig. 3: Cryo-EM reconstruction quality analysis of apo-ASK1–MAX2. A.** Two orientations of the atomic model of apo-ASK1–MAX2 (**PDB 9KLL**). ASK1 and MAX2 are coloured yellow and green, respectively. The F-box and LRR numbering is labelled. **B.** Cryo-EM density coloured by local resolution. Higher resolutions are coloured blue and lower resolutions are coloured red. **C.** Fourier shell correlation (FSC) curves for the cryo-EM reconstructions. Black line represents the correlation between independently processed half-maps, with resolution determined at FSC = 0.143. Blue line shows the correlation between the atomic model and the full cryo-EM map, with resolution assessed at FSC = 0.5. **D.** Euler angle distribution of the final reconstruction.

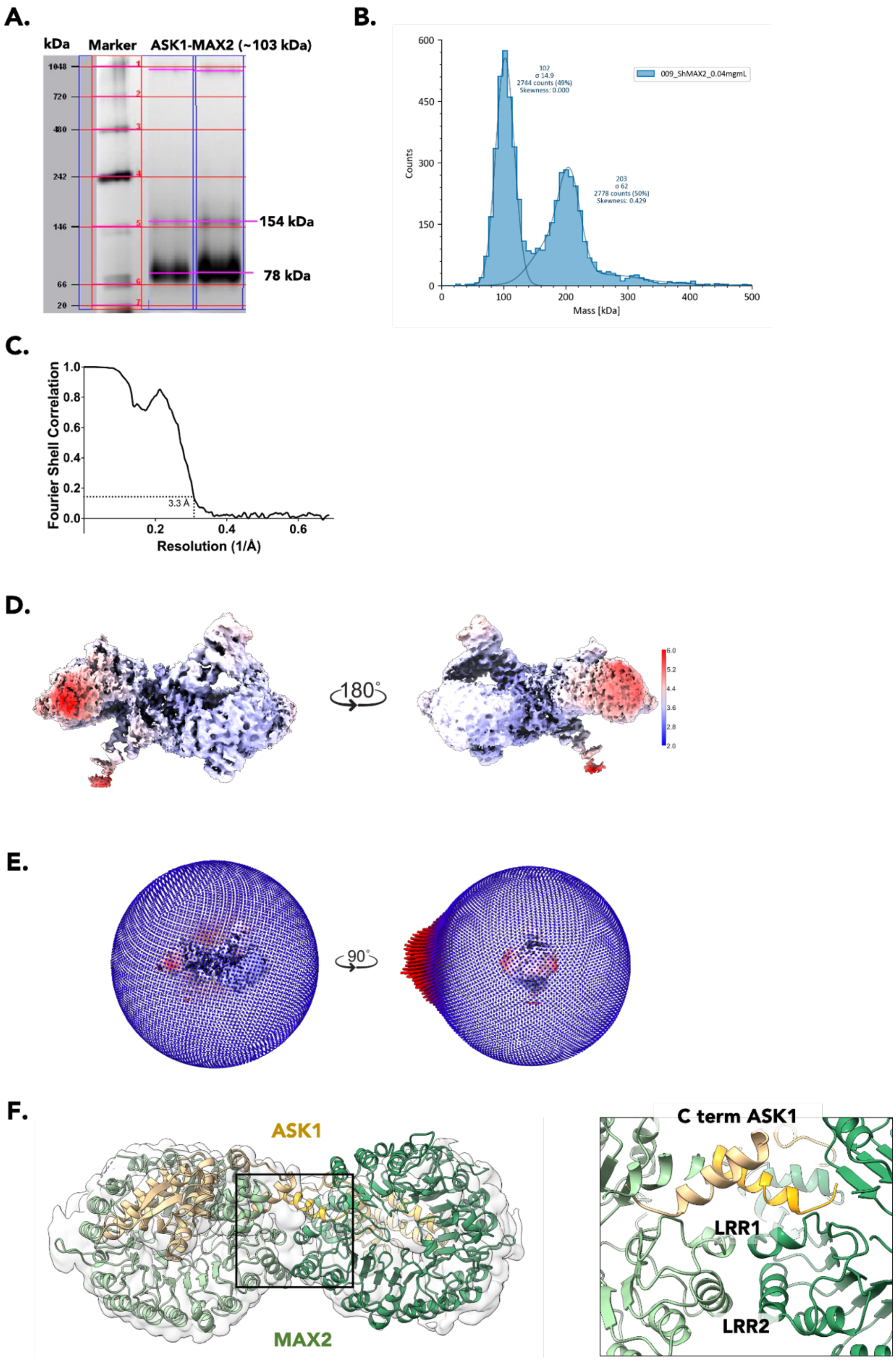
A.
kDa
Marker
ASK1-MAX2 (~103 kDa)
1048
720
480
242
146
66
20
154 kDa
78 kDa
B.
102
σ 14.9
2744 counts (49%)
Skewness: 0.000
203
σ 62
2778 counts (50%)
Skewness: 0.429
009_ShMAX2_0.04mgmL
Counts
Mass [kDa]
C.
Fourier Shell Correlation
3.3 Å
Resolution (1/Å)
D.
180°
E.
90°
F.
ASK1
MAX2
C term ASK1
LRR1
LRR2

**Supplementary Fig. 4: Biochemical characterisation and cryo-EM reconstruction quality analysis of the 2:2 ASK1–MAX2 dimer. A.** Native-PAGE of purified ASK1–MAX2 at low (10 µg; left) and high (20 µg; right) loading. Molecular weight was analysed using the ImageLab software from the Gel Doc EZimager (BioRad), using the NativeMark (TFS) marker migration distance as a reference. The ratio between the migration of the two strong bands is consistent with a dimeric species. Uncropped gel shown in Supplementary Fig. 13.**B.** Mass photometry histogram showing the mass distribution of 0.39 µM of ASK1–MAX2. The expected mass is 103 kDa for the monomer, while the estimated mass shows two distributions at 102 ± 15 kDa and 203 ± 62 kDa. **C.** Fourier shell correlation (FSC) curves for the cryo-EM reconstructions. Black line represents the correlation between independently processed half-maps, with resolution determined at FSC = 0.143. **D.** Cryo-EM density coloured by local resolution. Higher resolutions are coloured blue and lower resolutions are coloured red. **E.** Euler angle distribution of the final reconstruction. **F.** ASK1–MAX2 dimer interface. The atomic model fit in the transparent cryo-EM density (left). Black box outlines the magnified image (right) showing the dimer interface details. ASK1 and MAX2 are coloured yellow and green, respectively.

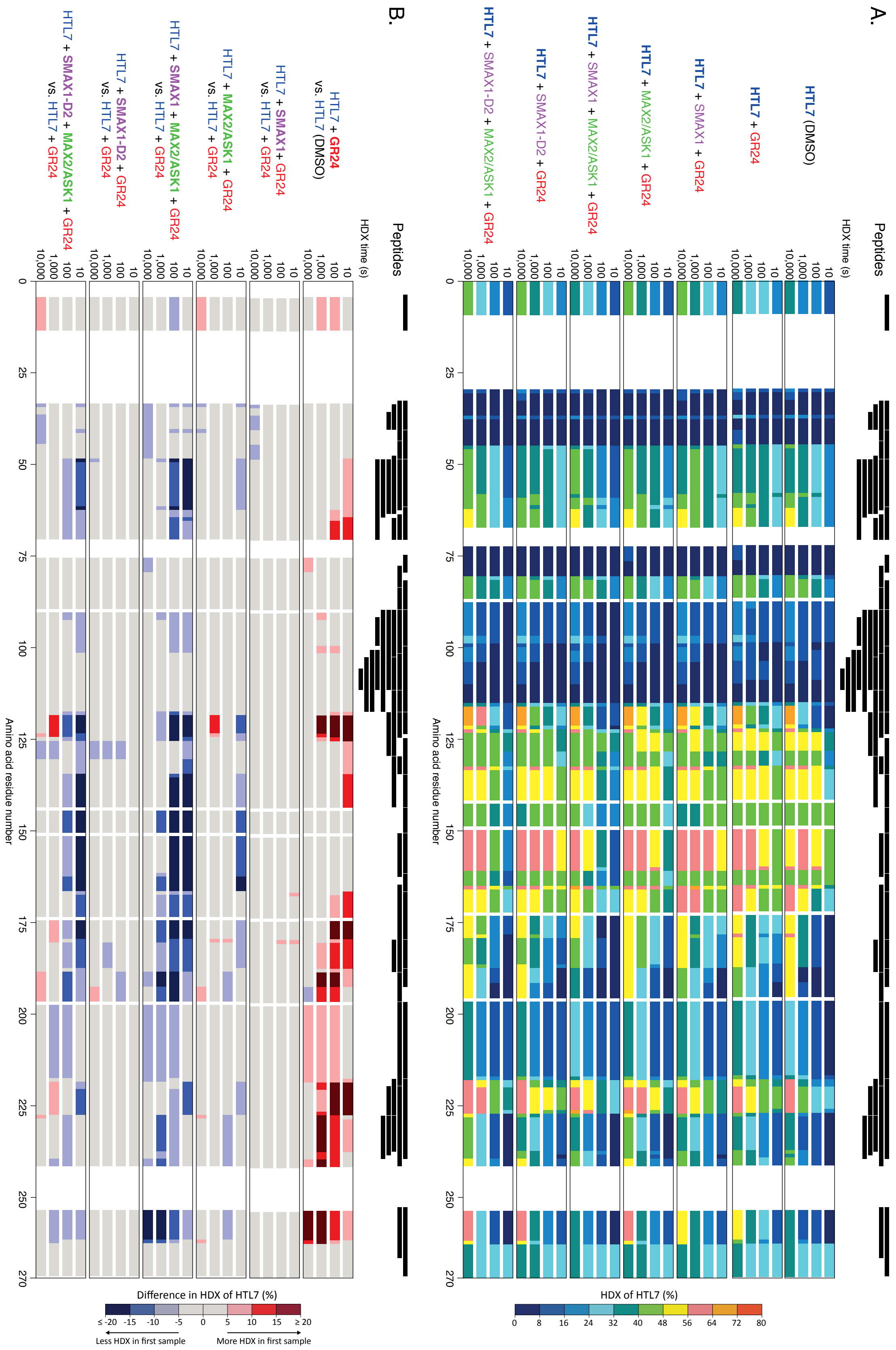

A.
Peptides
HDX time (s)
10
100
1,000
10,000
HTL7 (DMSO)
HTL7 + GR24
HTL7 + SMAX1 + GR24
HTL7 + MAX2/ASK1 + GR24
HTL7 + SMAX1 + MAX2/ASK1 + GR24
HTL7 + SMAX1-D2 + GR24
HTL7 + SMAX1-D2 + MAX2/ASK1 + GR24
0
25
50
75
100
125
150
175
200
225
250
270
Amino acid residue number
HDX of HTL7 (%)
0
8
16
24
32
40
48
56
64
72
80
B.
Peptides
HDX time (s)
HTL7 + GR24 vs. HTL7 (DMSO)
HTL7 + SMAX1 + GR24 vs. HTL7 + GR24
HTL7 + MAX2/ASK1 + GR24 vs. HTL7 + GR24
HTL7 + SMAX1 + MAX2/ASK1 + GR24 vs. HTL7 + GR24
HTL7 + SMAX1-D2 + GR24 vs. HTL7 + GR24
HTL7 + SMAX1-D2 + MAX2/ASK1 + GR24 vs. HTL7 + GR24
Amino acid residue number
Difference in HDX of HTL7 (%)
≤ -20
-15
-10
-5
0
5
10
15
≥ 20
Less HDX in first sample
More HDX in first sample

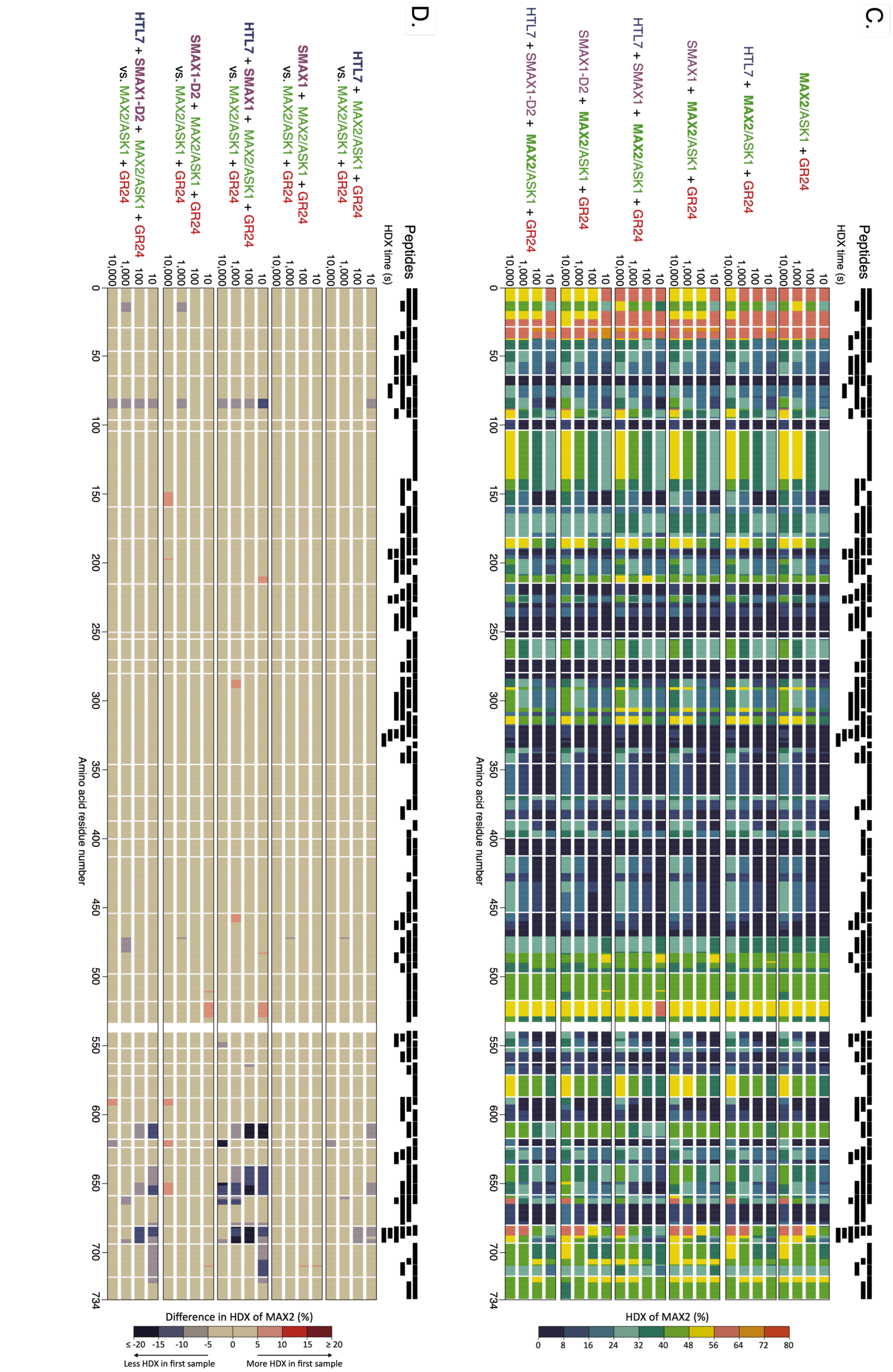


**Supplementary Fig. 5: HDX-MS analysis of HTL7 and MAX2. A.** Each black bar represents a peptide of HTL7 that was identified in HDX-MS experiments. The HDX per residue and deuteration time-point is plotted on the HTL7 amino acid sequence. **B.** The difference in HDX of HTL7 of pairwise comparisons between HTL7-containing samples is displayed on its amino acid sequence. Different tones of red and blue indicate HTL7 residues that incorporate more or less deuterium, respectively, in the first state of the pairwise comparison. Numerical values for deuterium uptake by peptides and residue-specific HDX are provided in **Supplementary Dataset 1. C.** Each black bar represents a peptide of MAX2 that was identified in HDX-MS experiments. The HDX per residue and deuteration time-point is plotted on the MAX2 amino acid sequence. **D.** The difference in HDX of MAX2 of pairwise comparisons between MAX2-containing samples is displayed on its amino acid sequence. Different tones of red and blue indicate MAX2 residues that incorporate more or less deuterium, respectively, in the first state of the pairwise comparison. Numerical values for deuterium uptake by peptides and residue-specific HDX are provided in **Supplementary Dataset 1**.

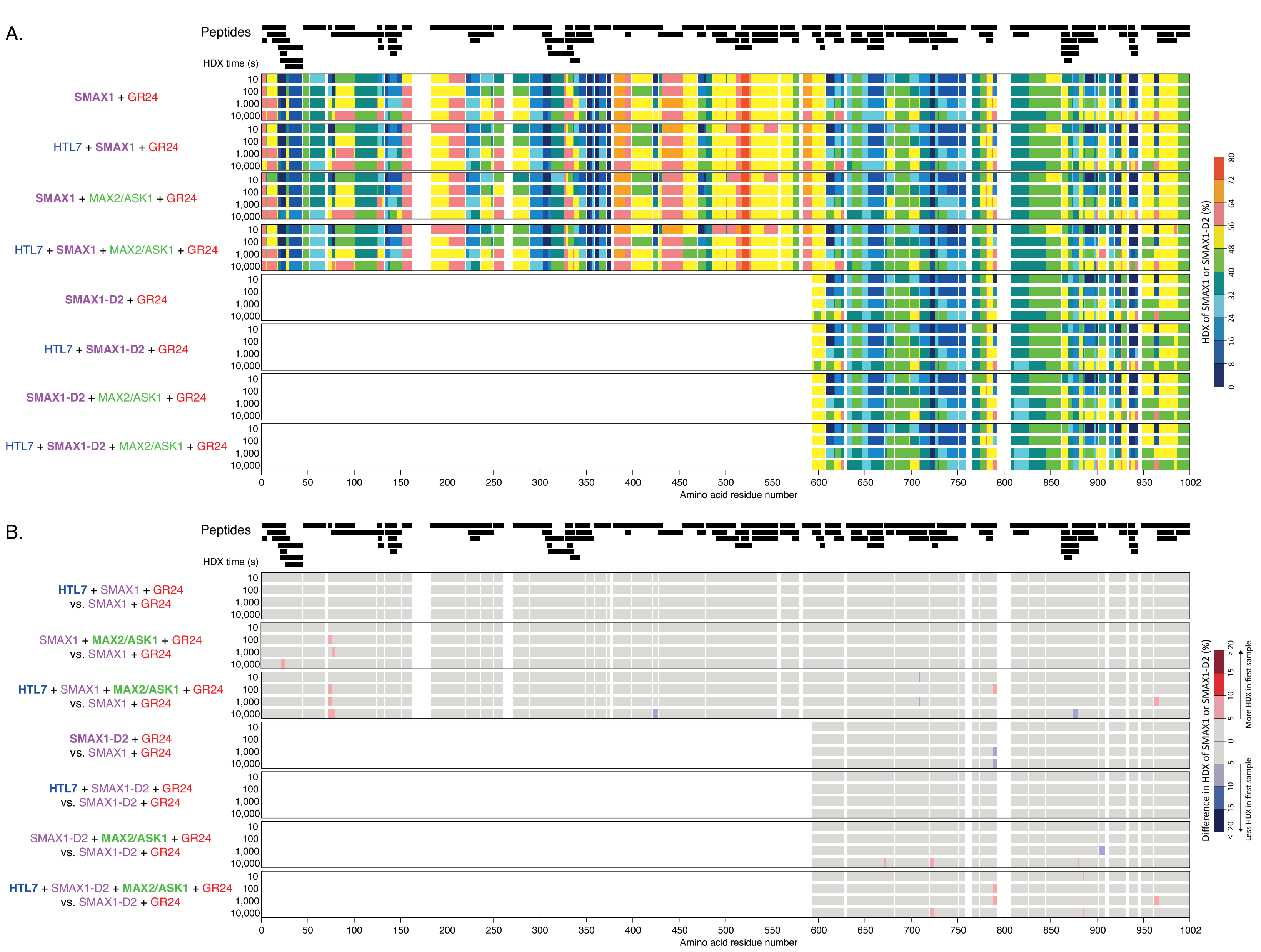
A.
Peptides
HDX time (s)
SMAX1 + GR24
HTL7 + SMAX1 + GR24
SMAX1 + MAX2/ASK1 + GR24
HTL7 + SMAX1 + MAX2/ASK1 + GR24
SMAX1-D2 + GR24
HTL7 + SMAX1-D2 + GR24
SMAX1-D2 + MAX2/ASK1 + GR24
HTL7 + SMAX1-D2 + MAX2/ASK1 + GR24
10
100
1,000
10,000
Amino acid residue number
HDX of SMAX1 or SMAX1-D2 (%)
B.
Peptides
HDX time (s)
HTL7 + SMAX1 + GR24 vs. SMAX1 + GR24
SMAX1 + MAX2/ASK1 + GR24 vs. SMAX1 + GR24
HTL7 + SMAX1 + MAX2/ASK1 + GR24 vs. SMAX1 + GR24
SMAX1-D2 + GR24 vs. SMAX1 + GR24
HTL7 + SMAX1-D2 + GR24 vs. SMAX1-D2 + GR24
SMAX1-D2 + MAX2/ASK1 + GR24 vs. SMAX1-D2 + GR24
HTL7 + SMAX1-D2 + MAX2/ASK1 + GR24 vs. SMAX1-D2 + GR24
Amino acid residue number
Difference in HDX of SMAX1 or SMAX1-D2 (%)
Less HDX in first sample
More HDX in first sample

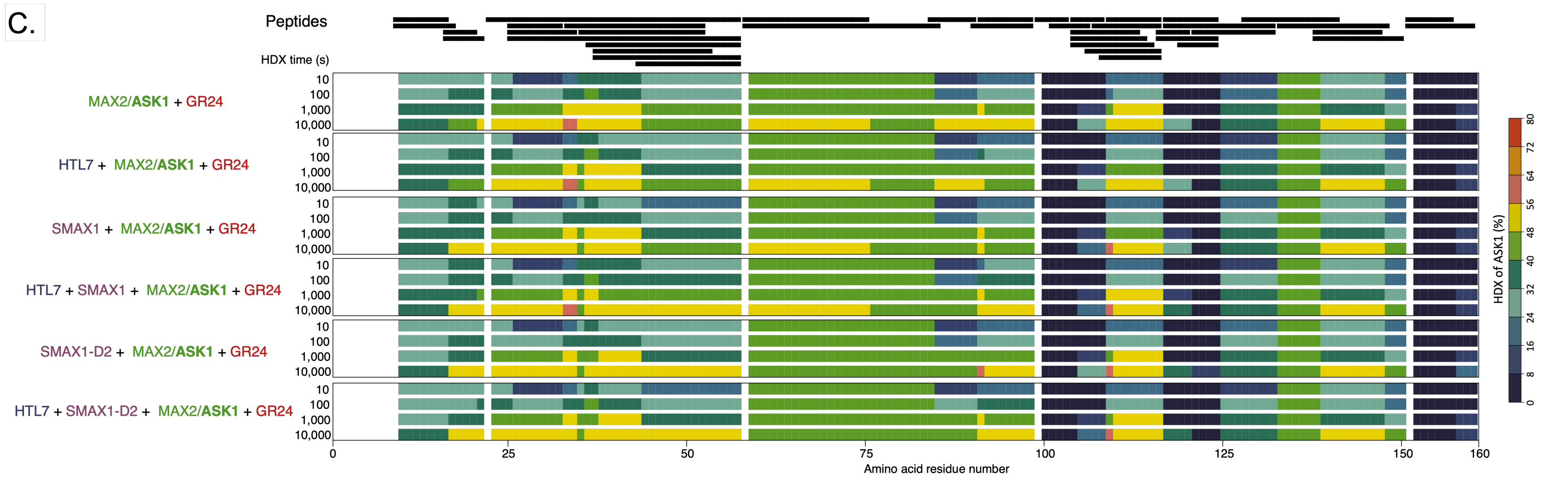
C.
Peptides
HDX time (s)
MAX2/ASK1 + GR24
HTL7 + MAX2/ASK1 + GR24
SMAX1 + MAX2/ASK1 + GR24
HTL7 + SMAX1 + MAX2/ASK1 + GR24
SMAX1-D2 + MAX2/ASK1 + GR24
HTL7 + SMAX1-D2 + MAX2/ASK1 + GR24
10
100
1,000
10,000
0
25
50
75
100
125
150
160
Amino acid residue number
HDX of ASK1 (%)
0
8
16
24
32
40
48
56
64
72
80

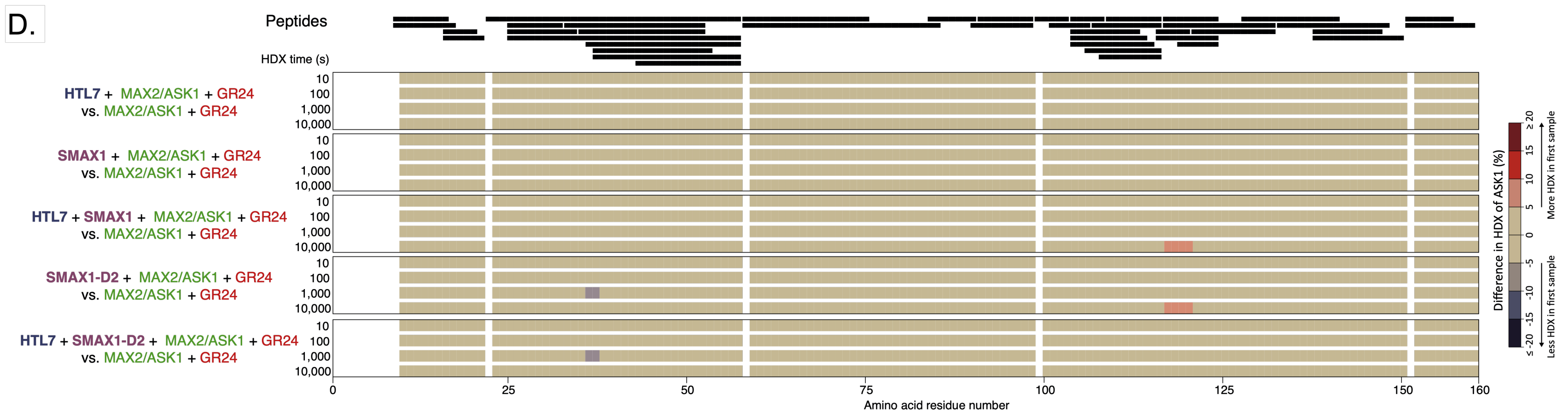
D.
Peptides
HDX time (s)
HTL7 + MAX2/ASK1 + GR24
vs. MAX2/ASK1 + GR24
SMAX1 + MAX2/ASK1 + GR24
vs. MAX2/ASK1 + GR24
HTL7 + SMAX1 + MAX2/ASK1 + GR24
vs. MAX2/ASK1 + GR24
SMAX1-D2 + MAX2/ASK1 + GR24
vs. MAX2/ASK1 + GR24
HTL7 + SMAX1-D2 + MAX2/ASK1 + GR24
vs. MAX2/ASK1 + GR24
10
100
1,000
10,000
0
25
50
75
100
125
150
160
Amino acid residue number
Difference in HDX of ASK1 (%)
≤ -20
-15
-10
-5
0
5
10
15
≥ 20
Less HDX in first sample
More HDX in first sample

**Supplementary Fig. 6: HDX-MS analysis of SMAX1 and ASK1. A.** Each black bar represents a peptide of SMAX1 that was identified in HDX-MS experiments. The HDX per residue and deuteration time-point is plotted on the SMAX1 amino acid sequence. **B.** The difference in HDX of SMAX1 of pairwise comparisons between SMAX1-containing samples is displayed on its amino acid sequence. Different tones of red and blue indicate SMAX1 residues that incorporate more or less deuterium, respectively, in the first state of the pairwise comparison. Numerical values for deuterium uptake by peptides and residue-specific HDX are provided in **Supplementary Dataset 1**. **C.** Each black bar represents a peptide of ASK1 that was identified in HDX-MS experiments. The HDX per residue and deuteration time-point is plotted on the ASK1 amino acid sequence. **D.** The difference in HDX of ASK1 of pairwise comparisons between ASK1-containing samples is displayed on its amino acid sequence. Different tones of red and blue indicate ASK1 residues that incorporate more or less deuterium, respectively, in the first state of the pairwise comparison. Numerical values for deuterium uptake by peptides and residue-specific HDX are provided in **Supplementary Dataset 1.**

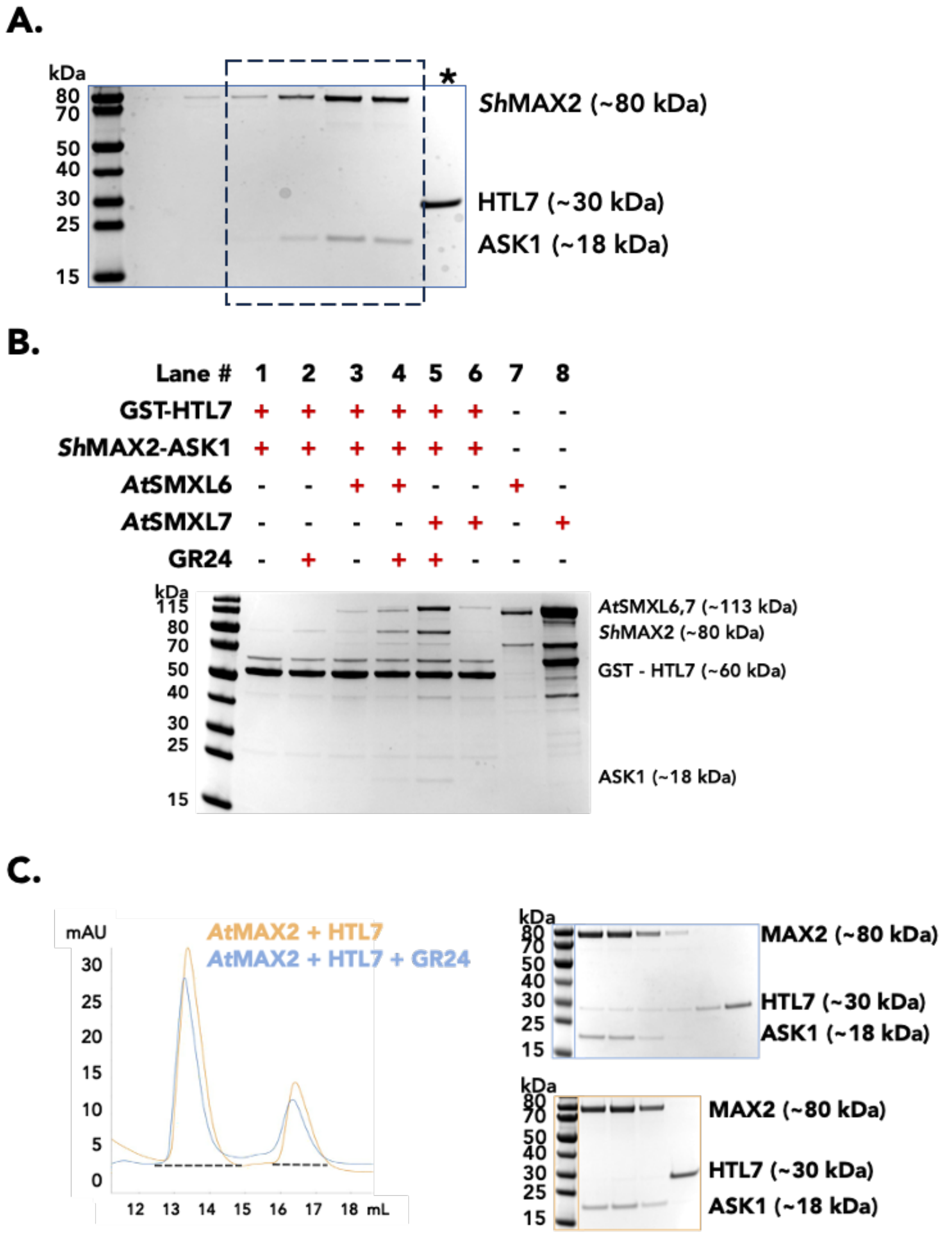


**Supplementary Fig. 7: Biochemical support for MAX2–HTL7–SMAX1$_{D2}$ interactions.**
**A.** Corresponding to **Fig. 4G**. Coomassie stained SDS-PAGE of MAX2–HTL7 and GR24 alone. Asterisk and dashed box show the fractions analysed from the SEC profile in Fig. 4G. **B.** Coomassie stained SDS-PAGE of GST pull-down assays using GST-HTL7 as a bait and combinations of ASK1–MAX2, SMXL6, SMXL7, and GR24. Plus signs denote presence and minus signs absence of species in each pull-down assay. Approximate molecular weights are labelled. **C.** SEC profiles (left) and corresponding Coomassie stained SDS-PAGE (right) of ASK1–*At*MAX2–HTL7 with (blue) and without (orange) GR24. Solid lines show the absorbance at 280 nm. Black dashed lines highlight the fractions analysed by SDS-PAGE. The corresponding SDS-PAGE gel for each SEC profile is outlined with the same colour. Approximate molecular weights are labelled. Uncropped gels shown in Supplementary Fig. 13.

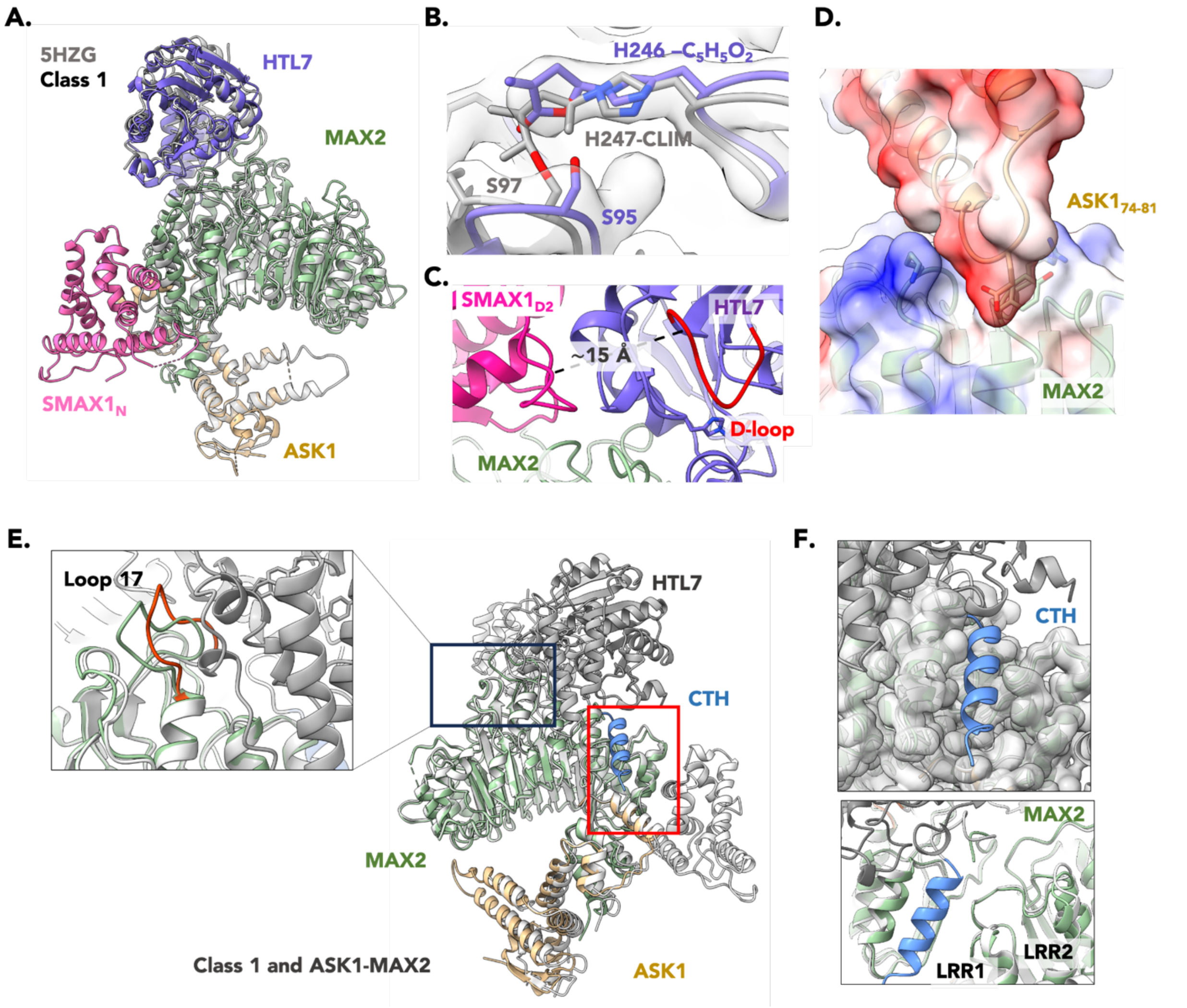


**Supplementary Fig. 8: Comparison of 5HZG and Class 1 and apo vs. HTL7-bound structures. A.** Superimposition of our cryo-EM structure of Class 1 (**PDB 9KKX**; coloured as in Fig. 1) and PDB 5HZG (white). **B.** Comparison of HTL7 from Class 1 (purple) with *At*D14 from PDB 5HZG (white). Cryo-EM density of Class 1 is shown as a transparent surface. The D-ring (our model) and CLIM modifications (proposed in PDB 5HZG) and the catalytic His and Ser residues are shown as sticks and labelled. **C.** Position of SMAX1$_{D2}$ relative to the D-loop of HTL7. Coloured as in Fig. 1. The D-loop is coloured red. The distance of ~15 Å is marked as a dotted line. **D.** Interface of ASK1–MAX2 through the tip of an extended helix-turn-helix element of ASK1 (residues 74–81) to the MAX2 LRR11 and LRR12. Interacting residues are shown as sticks. Transparent surfaces of ASK1 and MAX2 are shown and coloured by electrostatic potential, from red (negatively charged) to blue (positively charged). **E.** Superimposition of Class 1 (grey) with apo-ASK1–MAX2 (coloured) atomic models highlighting the engaged state of ASK1 in Class 1, relative to the resting state in apo-ASK1–MAX2. The black box magnifies the MAX2 LRR17 loop repositioning in the HTL7-bound Class 1 (red), relative to apo-ASK1–MAX2. The MAX2 CTH from Class 1 is coloured blue. Red box corresponds to panel F. **F.** Magnification of the red boxed region from panel E. Top panel shows the absence of cryo-EM density for the MAX2 CTH in the apo-ASK1–MAX2 reconstruction. The

Class 1 atomic model is overlaid on the apo-ASK1–MAX2 cryo-EM density. Bottom panel shows the shift of MAX2 LRR1 and LRR2 when the CTH is present in Class 1 relative to apo-ASK1–MAX2. Coloured as in E.

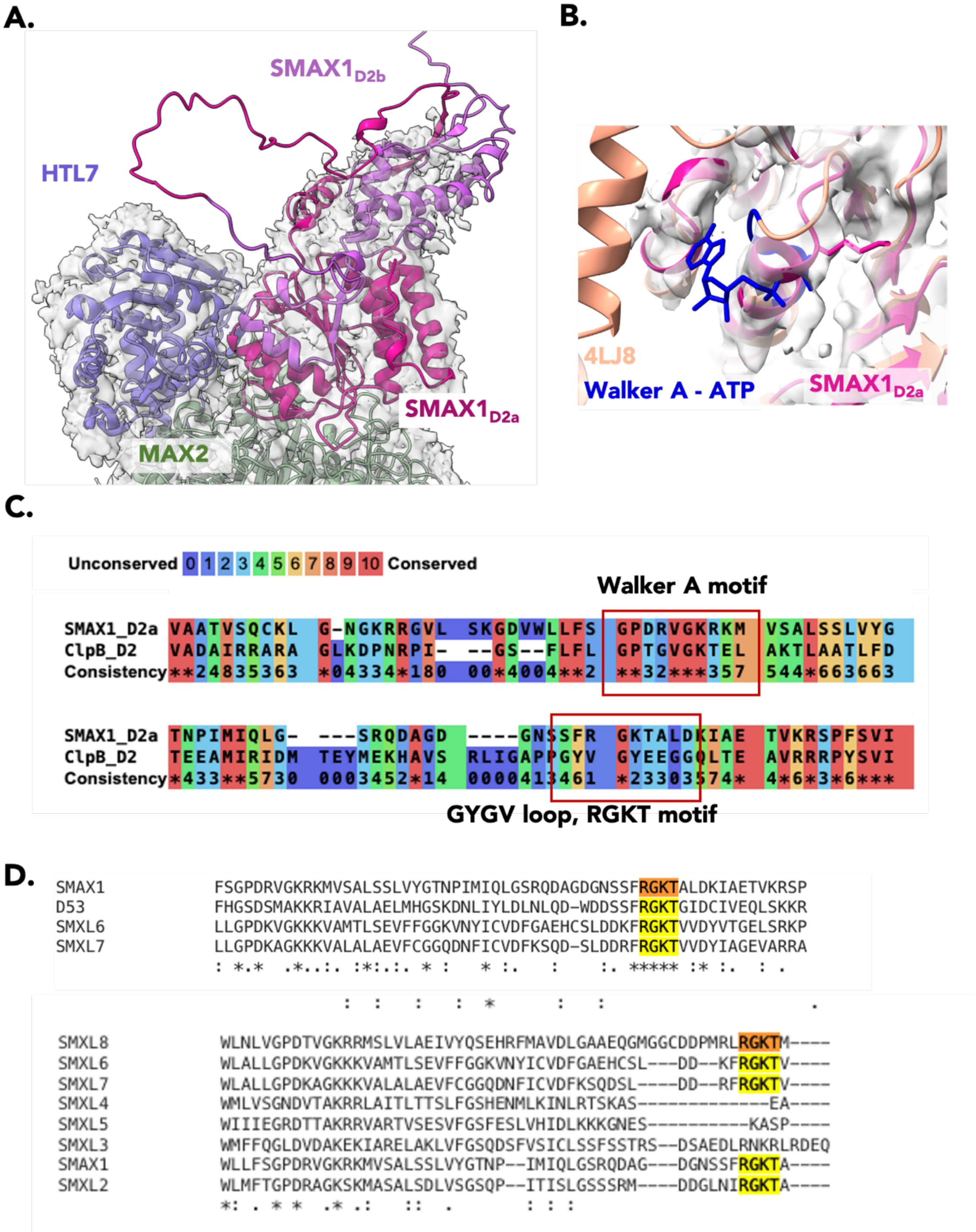


**Supplementary Fig. 9: Analysis of SMAX1$_{D2}$. A.** Cryo-EM density of Class 2 displayed at low threshold with a model of full-length SMAX1$_{D2}$ overlaid. MAX2 and HTL7 are coloured green and purple, respectively. SMAX1$_{D2a}$ and SMAX1$_{D2b}$ are coloured dark pink and light purple, respectively. **B.** Superimposition of SMAX1$_{D2}$ from Class 2 (dark pink) and PDB 4LJ8 (orange). The Walker A fold and ATP of PDB 4LJ8 are coloured blue. Cryo-EM density of Class 2 at low threshold is shown as a transparent grey surface. **C.** Sequence conservation of the Walker A motif (top) and pore loop (bottom) between $SMAX1_{D2}$ and ClpB. Sequences aligned using the PRALINE program. Red amino acids show high conservation and blue low conservation. Boxes highlight the respective motifs. **D.** Sequence alignment of the $SMAX1_{D2}$ RGKT motif across rice and Arabidopsis homologues. The RGKT motif is highlighted. Sequences aligned using MUSCLE.

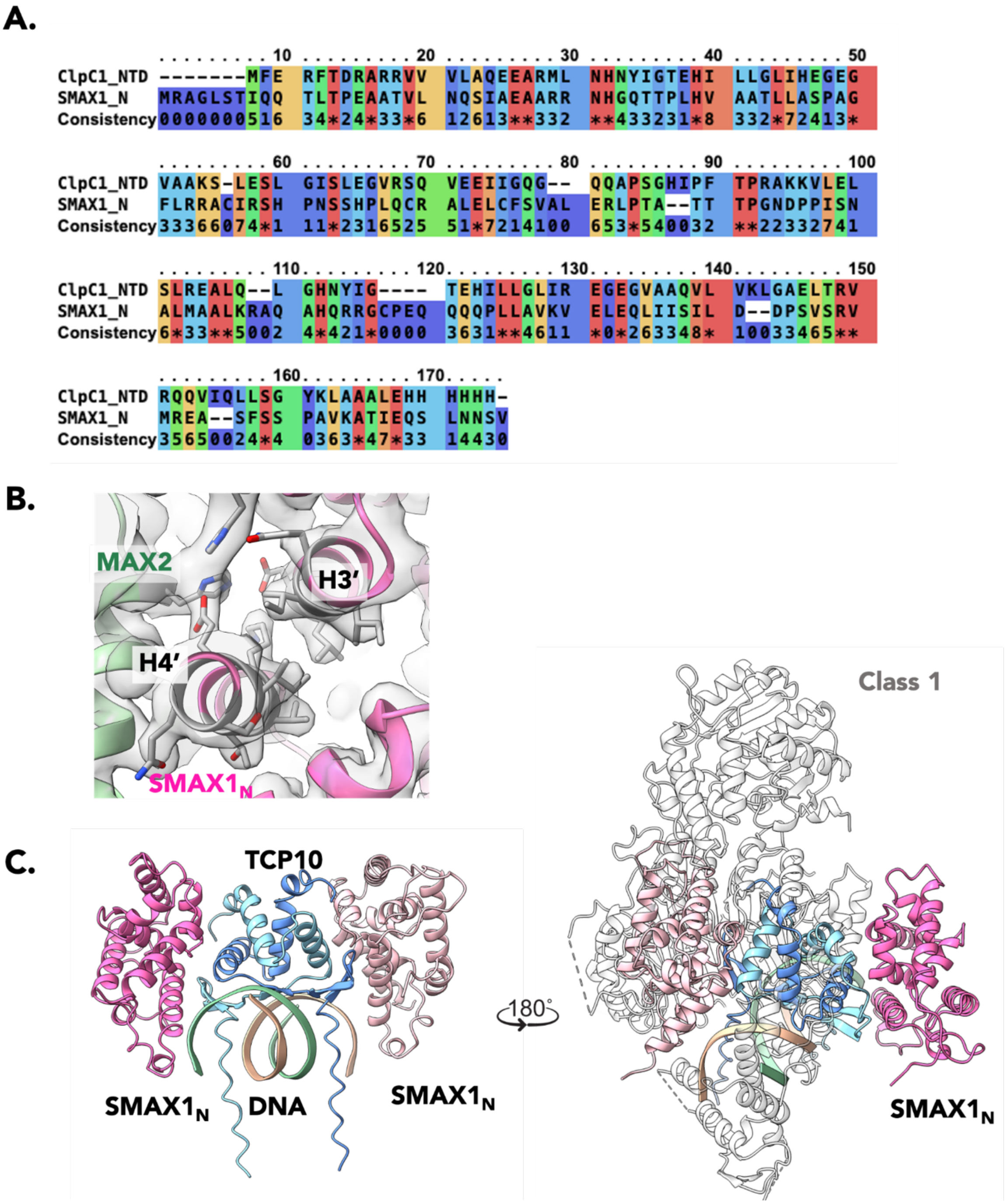


**Supplementary Fig. 10: Analysis of the SMAX1$_N$ interface. A.** Sequence alignment of SMAX1$_N$ with the NTD of ClpC1 (PDB 6PBA). Red amino acids show high conservation and blue low conservation. **B.** Fit of SMAX1$_N$ in the cryo-EM density of Class 1 (transparent grey surface). Helices are labelled and side chains are shown as grey sticks. MAX2 is green and SMAX1$_N$ is light pink. **C.** (Left) AlphaFold-predicted model of the TCP10 dimer (blue) interacting with SMAX1$_N$ (pink) and DNA (cartoon ribbons). (Right) The AlphaFold TCP10–SMAX1$_N$–DNA model is superimposed on Class 1 showing the overlapping binding surfaces of SMAX1$_N$ with MAX2 or TCP10.

**A.**

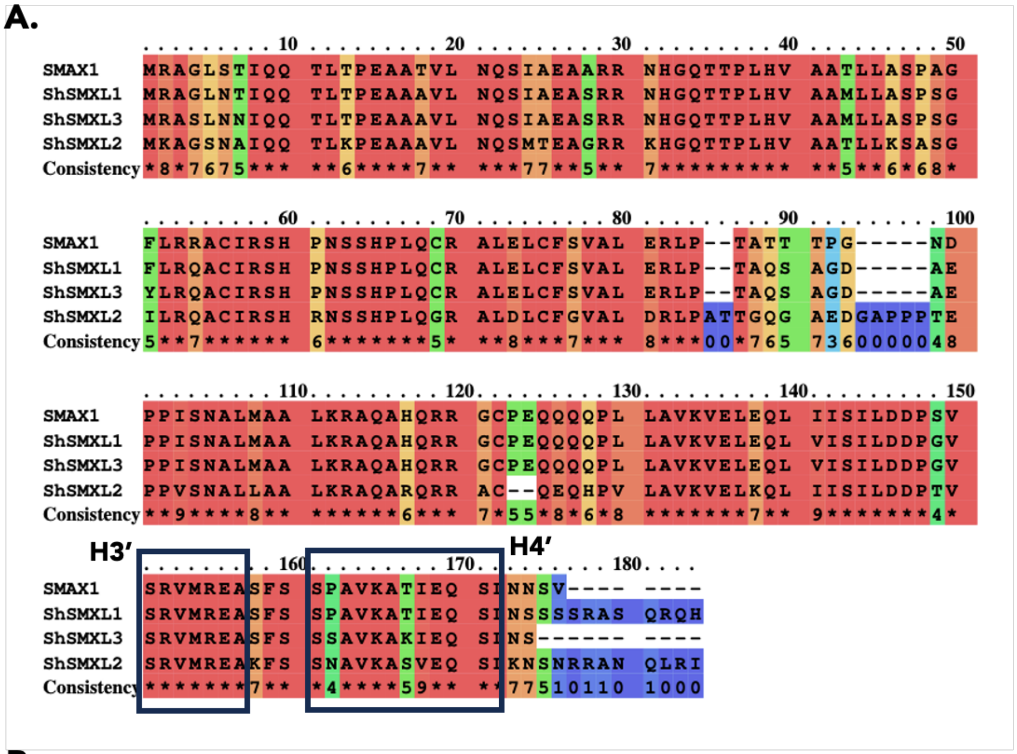


**B.**

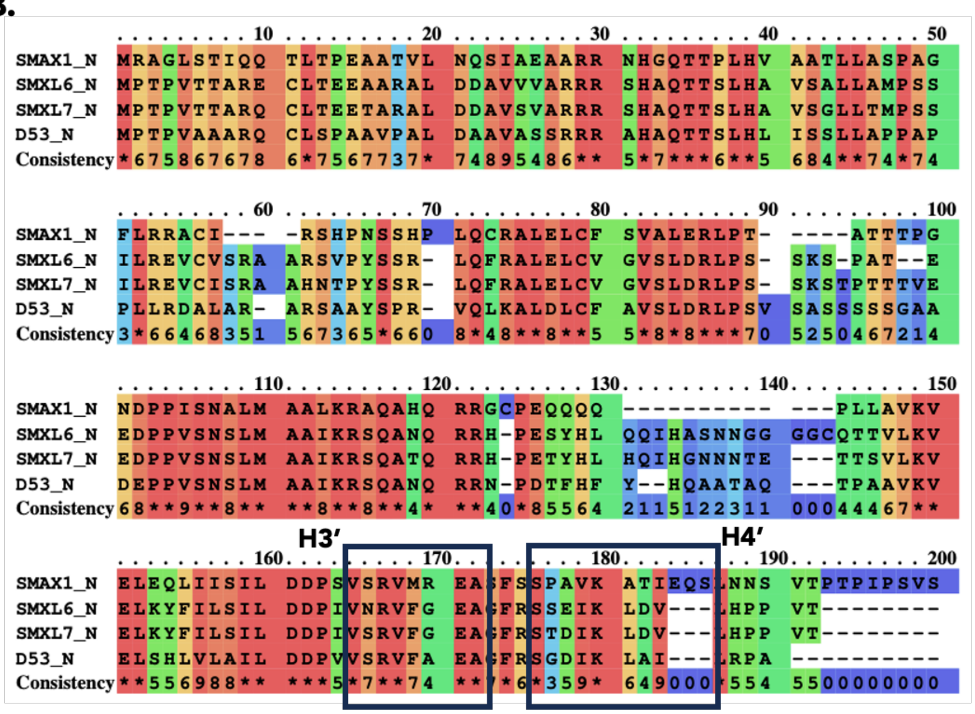


**C.**

| | | | | | | | |
|---|---|---|---|---|---|---|---|
| 1: ShSMXL2_N | 100.00 | 74.25 | 73.71 | 75.15 | 44.31 | 47.27 | 43.82 |
| 2: AtSMAX1_N | 74.25 | 100.00 | 89.94 | 87.43 | 44.51 | 46.99 | 47.02 |
| 3: ShSMXL1_N | 73.71 | 89.94 | 100.00 | 97.01 | 44.51 | 46.99 | 43.75 |
| 4: ShSMXL3_N | 75.15 | 87.43 | 97.01 | 100.00 | 44.51 | 46.99 | 45.18 |
| 5: OsD53_N | 44.31 | 44.51 | 44.51 | 44.51 | 100.00 | 57.14 | 55.62 |
| 6: AtSMXL6_N | 47.27 | 46.99 | 46.99 | 46.99 | 57.14 | 100.00 | 86.67 |
| 7: AtSMXL7_N | 43.82 | 47.02 | 43.75 | 45.18 | 55.62 | 86.67 | 100.00 |

**Supplementary Fig. 11: Sequence analysis of SMAX1$_N$. A.** Sequence alignment of SMAX1$_N$ with *Sh*SMXL1,2,3. The MAX2-interacting helices, H3' and H4', are boxed. Red amino acids show high conservation and blue low conservation. **B.** Sequence alignment of SMAX1$_N$ domain with *At*SMXL6,7 and *Os*D53. The MAX2-interacting helices, H3' and H4', are boxed. Red amino acids show high conservation and blue low conservation. **C.** Percent identity matrix (Clustal2.1) for all the SMAX1$_N$ homologues in A-B. The highest identity pairs are boxed. *At*SMAX1$_N$ shows highest identity to homologues in Striga.

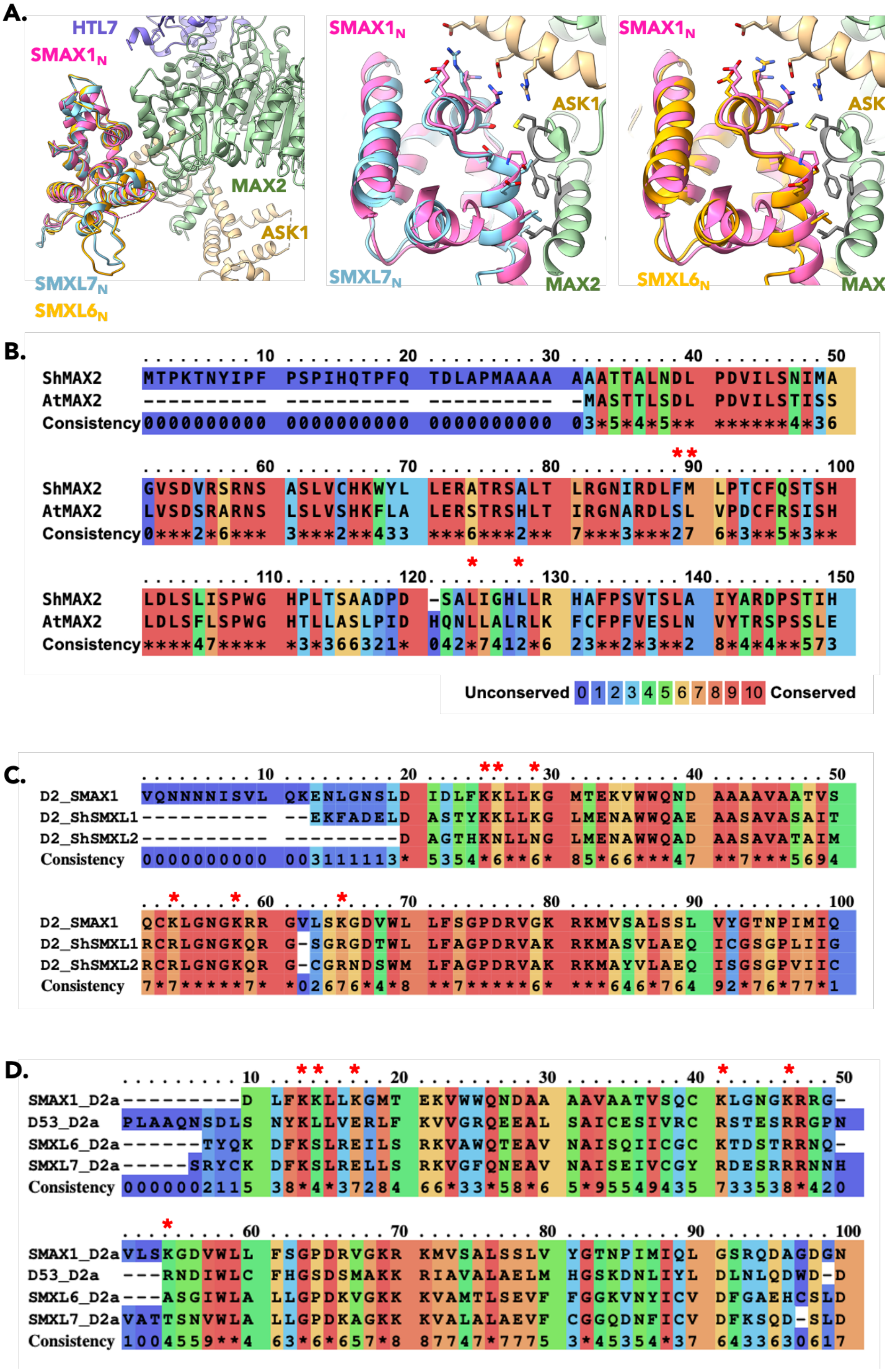


**Supplementary Fig. 12: Similarity of Arabidopsis and Striga proteins. A.** Superimposition of Class 3 (coloured as in Fig. 1) and AlphaFold predicted models of *At*SMXL7$_N$ (blue; middle) and

*At*SMXL6$_N$ (orange; right). Interacting residues are shown as sticks. Interacting residues on MAX2 are coloured grey. **B.** Sequence conservation of *Sh*MAX2 and *At*MAX2. Red asterisks mark the MAX2 residues involved in the hydrophobic interactions with the N domain depicted in A. Red amino acids show high conservation and blue low conservation. **C.** Sequence conservation between *At*SMAX1$_{D2}$, *Sh*SMXL1$_{D2}$ and *Sh*SMXL2$_{D2}$. Red asterisks mark the favourably placed Lys residues as in Fig. 6E. **D.** Sequence conservation between *At*SMAX1$_{D2}$ and *Os*D53$_{D2}$, *At*SMXL6$_{D2}$ and *At*SMXL7$_{D2}$. Red asterisks mark the favourably placed Lys residues as in Fig. 6E.

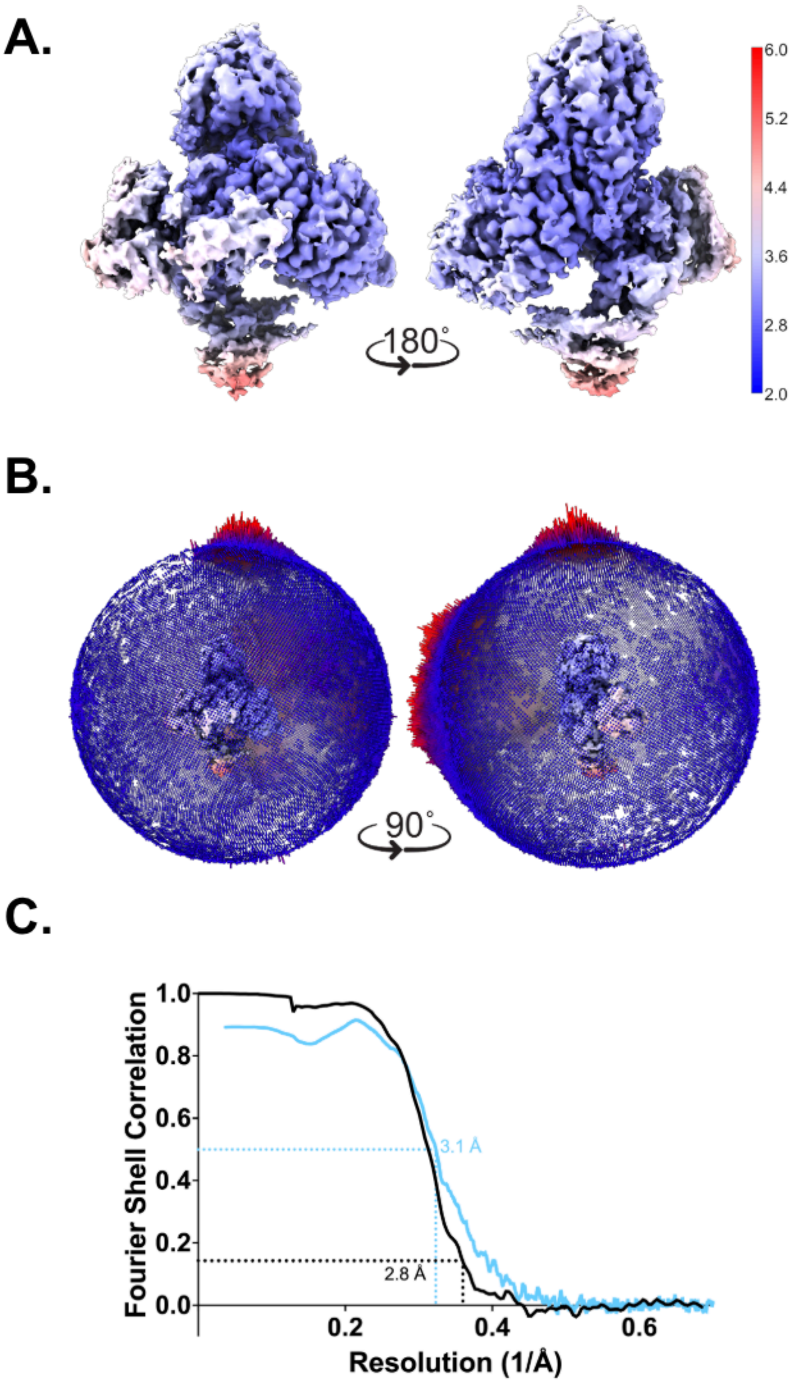


**Supplementary Fig. 13: Cryo-EM reconstruction quality analysis of ASK1–MAX2–HTL7–SMAX1 Class 1. A.** Cryo-EM density coloured by local resolution. Higher resolutions are coloured blue and lower resolutions are coloured red. **B.** Euler angle distribution of the final reconstruction.
**C.** Fourier shell correlation (FSC) curves for the cryo-EM reconstructions. Black line represents the correlation between independently processed half-maps, with resolution determined at FSC = 0.143. Blue line shows the correlation between the atomic model and the full cryo-EM map, with resolution assessed at FSC = 0.5.

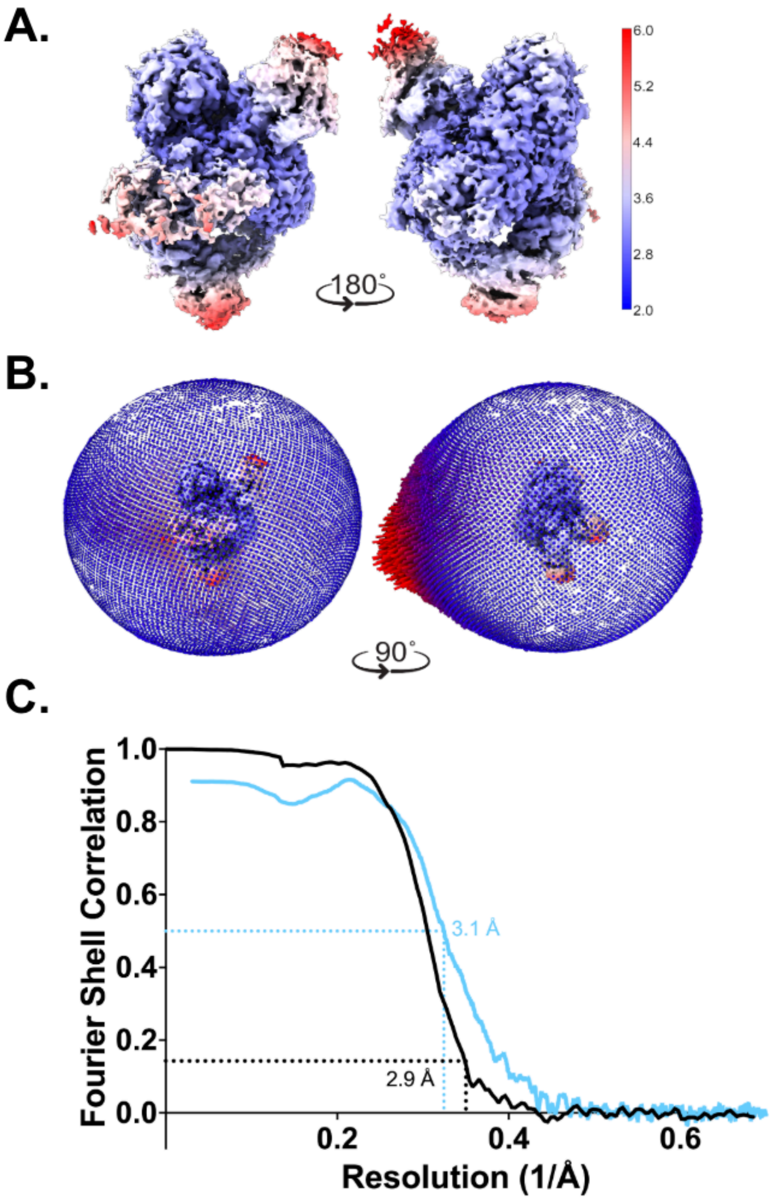


**Supplementary Fig. 14: Cryo-EM reconstruction quality analysis of ASK1–MAX2–HTL7–SMAX1 Class 2. A.** Cryo-EM density coloured by local resolution. Higher resolutions are coloured blue and lower resolutions are coloured red. **B.** Euler angle distribution of the final reconstruction. **C.** Fourier shell correlation (FSC) curves for the cryo-EM reconstructions. Black line represents the correlation between independently processed half-maps, with resolution determined at FSC = 0.143. Blue line shows the correlation between the atomic model and the full cryo-EM map, with resolution assessed at FSC = 0.5.

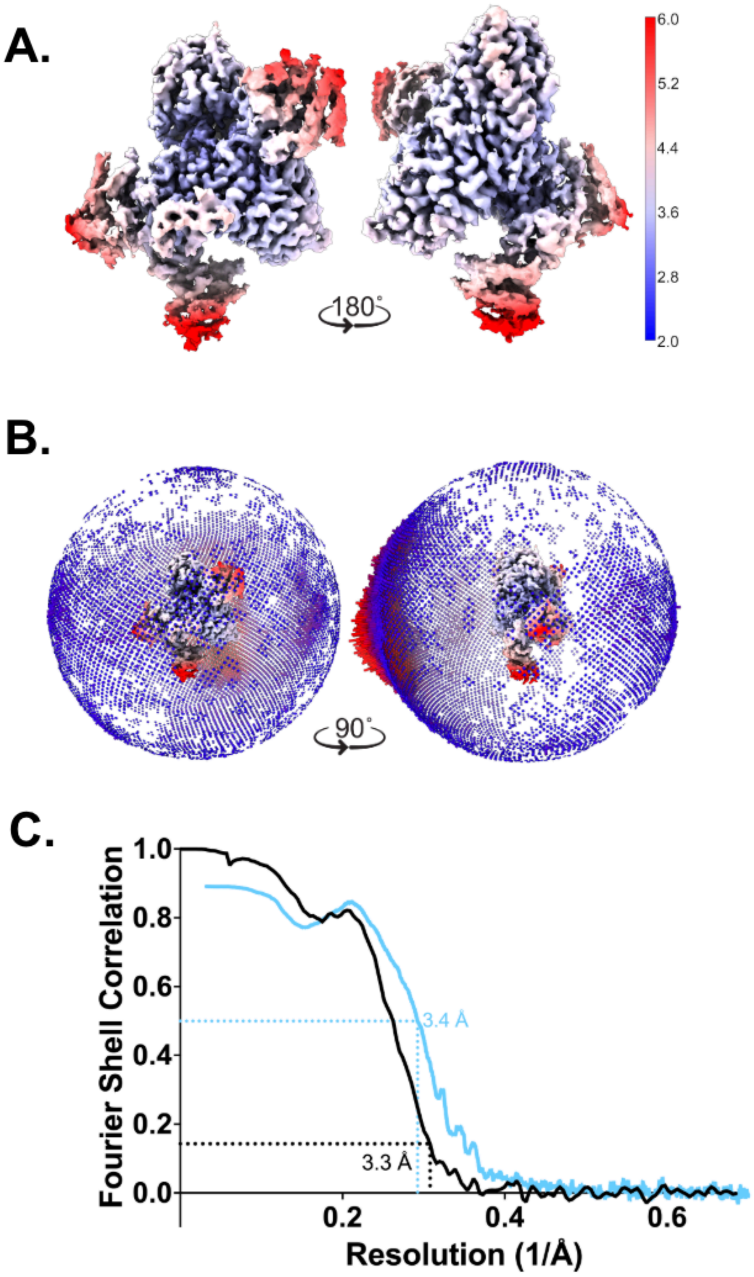


**Supplementary Fig. 15: Cryo-EM reconstruction quality analysis of ASK1–MAX2–HTL7–SMAX1 Class 3. A.** Cryo-EM density coloured by local resolution. Higher resolutions are coloured blue and lower resolutions are coloured red. **B.** Euler angle distribution of the final reconstruction. **C.** Fourier shell correlation (FSC) curves for the cryo-EM reconstructions. Black line represents the correlation between independently processed half-maps, with resolution determined at FSC = 0.143. Blue line shows the correlation between the atomic model and the full cryo-EM map, with resolution assessed at FSC = 0.5.

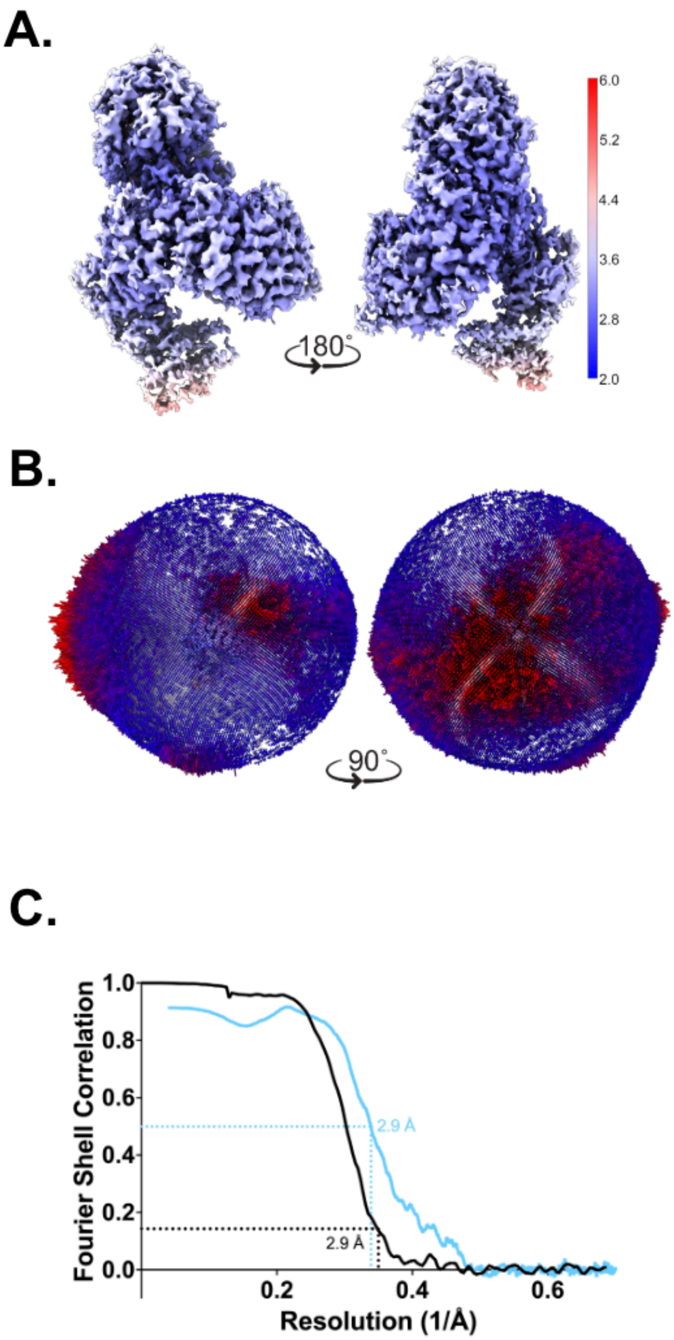


**Supplementary Fig. 16: Cryo-EM reconstruction quality analysis of ASK1–MAX2–HTL7–SMAX1 Class 4. A.** Cryo-EM density coloured by local resolution. Higher resolutions are coloured blue and lower resolutions are coloured red. **B.** Euler angle distribution of the final reconstruction. **C.** Fourier shell correlation (FSC) curves for the cryo-EM reconstructions. Black line represents the correlation between independently processed half-maps, with resolution determined at FSC = 0.143. Blue line shows the correlation between the atomic model and the full cryo-EM map, with resolution assessed at FSC = 0.5.

**Supplementary Figure 1**

**A.**

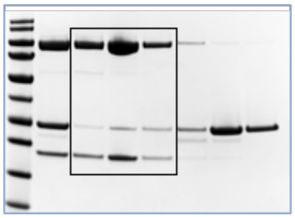

**C.**

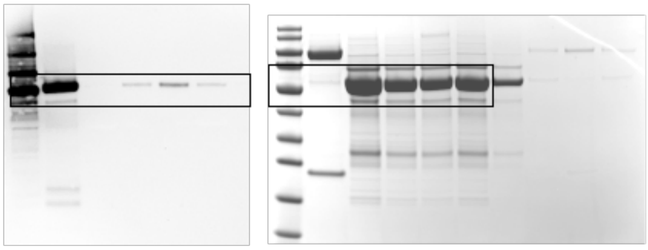

**Supplementary Figure 4**

**A.**

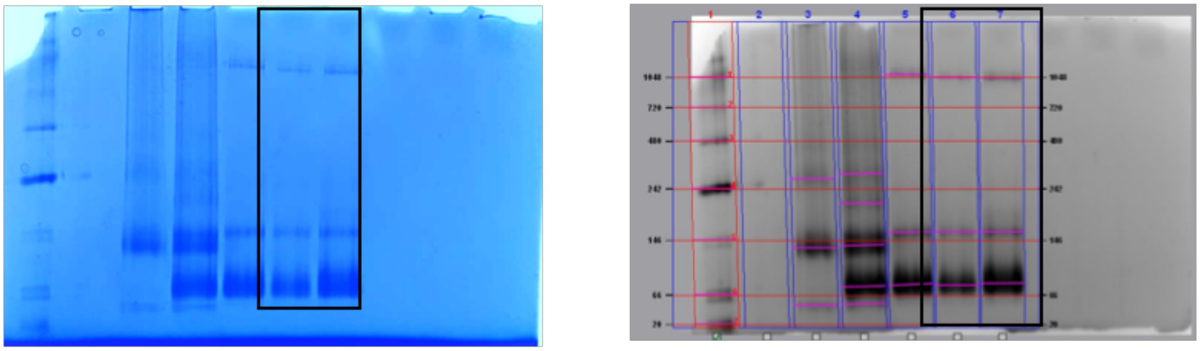

**Supplementary Figure 7**

**C.**

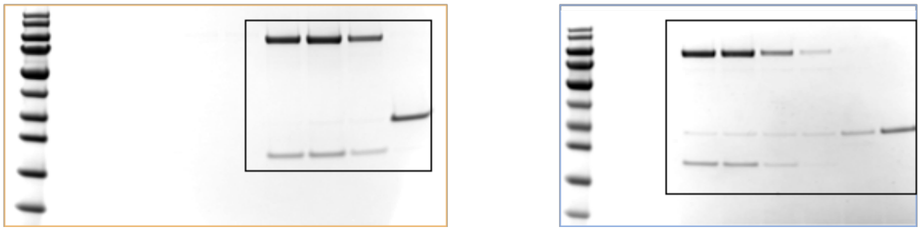

**Supplementary Fig. 17: Uncropped protein gels.**
